# The nucleotide messenger (p)ppGpp is a co-repressor of the purine synthesis transcription regulator PurR in Firmicutes

**DOI:** 10.1101/2020.12.02.409011

**Authors:** Brent W. Anderson, Maria A. Schumacher, Jin Yang, Asan Turdiev, Husan Turdiev, Qixiang He, Vincent T. Lee, Richard G. Brennan, Jue D. Wang

## Abstract

The nucleotide messenger (p)ppGpp allows bacteria to adapt to fluctuating environments by reprogramming the transcriptome. Yet despite its well-recognized role in gene regulation, (p)ppGpp is only known to directly affect transcription in Proteobacteria. Here we reveal a different mechanism of gene regulation by (p)ppGpp in Firmicutes from soil bacteria to pathogens: (p)ppGpp serves as a co-repressor of the transcription factor PurR to downregulate purine biosynthesis. We identified PurR as a receptor of (p)ppGpp in *Bacillus anthracis* and revealed that (p)ppGpp strongly enhances PurR binding to its regulon in the *Bacillus subtilis* genome. A co-structure reveals that (p)ppGpp binds to a PurR pocket reminiscent of the active site of PRT enzymes that has been repurposed to serve a purely regulatory role, where the effectors (p)ppGpp and PRPP compete to allosterically control transcription. PRPP inhibits PurR DNA binding to induce transcription of purine synthesis genes, whereas (p)ppGpp antagonizes PRPP to enhance PurR DNA binding and repress transcription. A (p)ppGpp-refractory *purR* mutant fails to downregulate purine synthesis genes upon starvation. Our work establishes precedent of (p)ppGpp as a classical transcription co-repressor and reveals the key function of (p)ppGpp in regulating nucleotide synthesis through gene regulation, from the human intestinal tract to host-pathogen interfaces.

## INTRODUCTION

Nucleotide second messengers allow organisms to rewire transcription to reprogram physiology and lifestyle for survival in changing environments. In bacteria, nucleotide second messengers, such as the cyclic nucleotides cAMP, c-di-AMP, and c-di-GMP, regulate transcription by binding to transcription factors as effectors and allosterically affecting the transcription factor’s binding to DNA regulatory elements (Fahmi et al., 2017; Gallagher et al., 2020; Hengge, 2009; Kalia et al., 2013; Sassone-Corsi, 1995; Tao et al., 2010; Tschowri et al., 2014; Zhang et al., 2013). In fact, cAMP, the key regulator of catabolite repression, was one of the first identified allosteric effectors of transcription, establishing the paradigm of gene regulation (Ullman et al., 1976).

In contrast to cyclic nucleotides, another group of signaling nucleotides is the linear ‘alarmone’ nucleotides, including the ubiquitous nucleotide messengers ppGpp and pppGpp, collectively (p)ppGpp. (p)ppGpp is necessary for bacteria to adapt to environmental changes from nutrient availability to antibiotic assault (Cashel et al., 1996; Kriel et al., 2012; Potrykus and Cashel, 2008). Since its discovery half a century ago, (p)ppGpp was best known for its profound role in gene regulation: downregulating the transcription of rRNA and tRNA operons and modulating global transcription in bacteria. (p)ppGpp binds RNA polymerase and the transcription factor DksA-RNA polymerase interface in Proteobacteria, and it binds the virulence regulator complex MglA-SspA in *Francisella tularensis* (Cuthbert et al., 2017; Ross et al., 2016). However, unlike cyclic nucleotide second messengers, (p)ppGpp has not been known to act as a canonical allosteric effector of a promoter-binding transcription factor in bacteria.

In addition, (p)ppGpp has only been shown to regulate transcription directly in Proteobacteria (Cuthbert et al., 2017; Ross et al., 2016; Sanchez-Vazquez et al., 2019). In other bacteria phyla, no transcription proteins have been shown to be directly regulated by (p)ppGpp. For example, in Firmicutes, including the soil bacterium *Bacillus subtilis* and the pathogen *Bacillus anthracis*, (p)ppGpp affects the transcription of hundreds of genes, but the only known mechanism is indirect: via control of levels of GTP, which is the initiating nucleotide for ribosomal RNA genes and is an effector for the global regulator CodY (Handke et al., 2008; Krásný and Gourse, 2004; Kriel et al., 2014).Therefore, the pervasiveness of (p)ppGpp’s role in direct gene expression remains unclear.

Here we report the first example of (p)ppGpp as a canonical allosteric effector of a transcription regulator, and the first example of transcription under direct control of (p)ppGpp in bacterial species beyond the Proteobacteria phylum. We found that (p)ppGpp directly regulates transcription in *Bacillus* by binding to the transcription regulator PurR and serving as its co-repressor. PurR controls expression of *de novo* purine nucleotide biosynthesis genes. (p)ppGpp induction, upon starvation, strongly increases PurR repression of *de novo* purine nucleotide biosynthesis. PurR proteins are highly conserved across Firmicutes, comprising a DNA binding domain and a regulatory domain resembling phosphoribosyltransferase (PRT) enzymes. PRT enzymes, including HPRT and XPRT, are purine salvage enzyme that uses the metabolite 5-phosphoribosyl-1-pyrophosphate (PRPP) to produce the purine nucleotides GMP, XMP and IMP. However, the PRT domain on PurR lacks enzymatic activity and serves a purely regulatory role by binding PRPP to de-repress transcription (Sinha et al., 2003; Weng et al., 1995). Structural analyses reveal that (p)ppGpp binds to the ‘active site’ of the PRT regulatory domain, where (p)ppGpp competes with PRPP to promote PurR binding to the promoter DNA, thus serving as a PurR co-repressor. Our study provides the first example of (p)ppGpp acting as a canonical allosteric effector of a transcription factor, thereby highlighting the conservation of signaling nucleotide as allosteric effectors of gene regulation, not only by cyclic but also linear signaling nucleotides.

## RESULTS

### Screening a *B. anthracis* ORF library identifies the purine transcription regulator PurR as a (p)ppGpp target

(p)ppGpp has been shown to bind to the RNA polymerase core enzyme in Proteobacteria (Molodtsov et al., 2018; Ross et al., 2013, 2016). In contrast, in Firmicutes (p)ppGpp regulates many targets, including nucleotide synthesis enzymes, ribosome associated GTPases, and DNA replication enzymes (Anderson et al., 2019, 2020; Corrigan et al., 2016; Kriel et al., 2012; Wang et al., 2007; Yang et al., 2020), but has not been shown to regulate transcription directly (Liu et al., 2015a). To uncover potential (p)ppGpp regulated transcriptional targets, we conducted a proteome based screen for proteins that directly bind (p)ppGpp in the pathogenic firmicute *Bacillus anthracis* (Yang et al., 2020). We screened the 5341 open reading frame (ORF) library from *B. anthracis*, placed each ORF into two expression constructs, one expressing the ORF with an N-terminal histidine (His) tag and the other with an N-terminal histidine maltose binding protein (HisMBP) tag. Each ORF in the His-tagged and HisMBP-tagged library was overexpressed and binding to ^32^P-pppGpp was assayed using differential radial capillary action of ligand assay (DRaCALA). The fraction ^32^P-pppGpp bound to protein in each lysate was normalized as a Z-score of each plate to reduce the influence of plate-to-plate variation. We used a strict Z-score cutoff of 2.5 to identify proteins very likely to be (p)ppGpp targets (Figure 1A-D), which yielded known and new (p)ppGpp targets (Table 1 and Table S1) (Yang et al., 2020).

**Figure 1.**
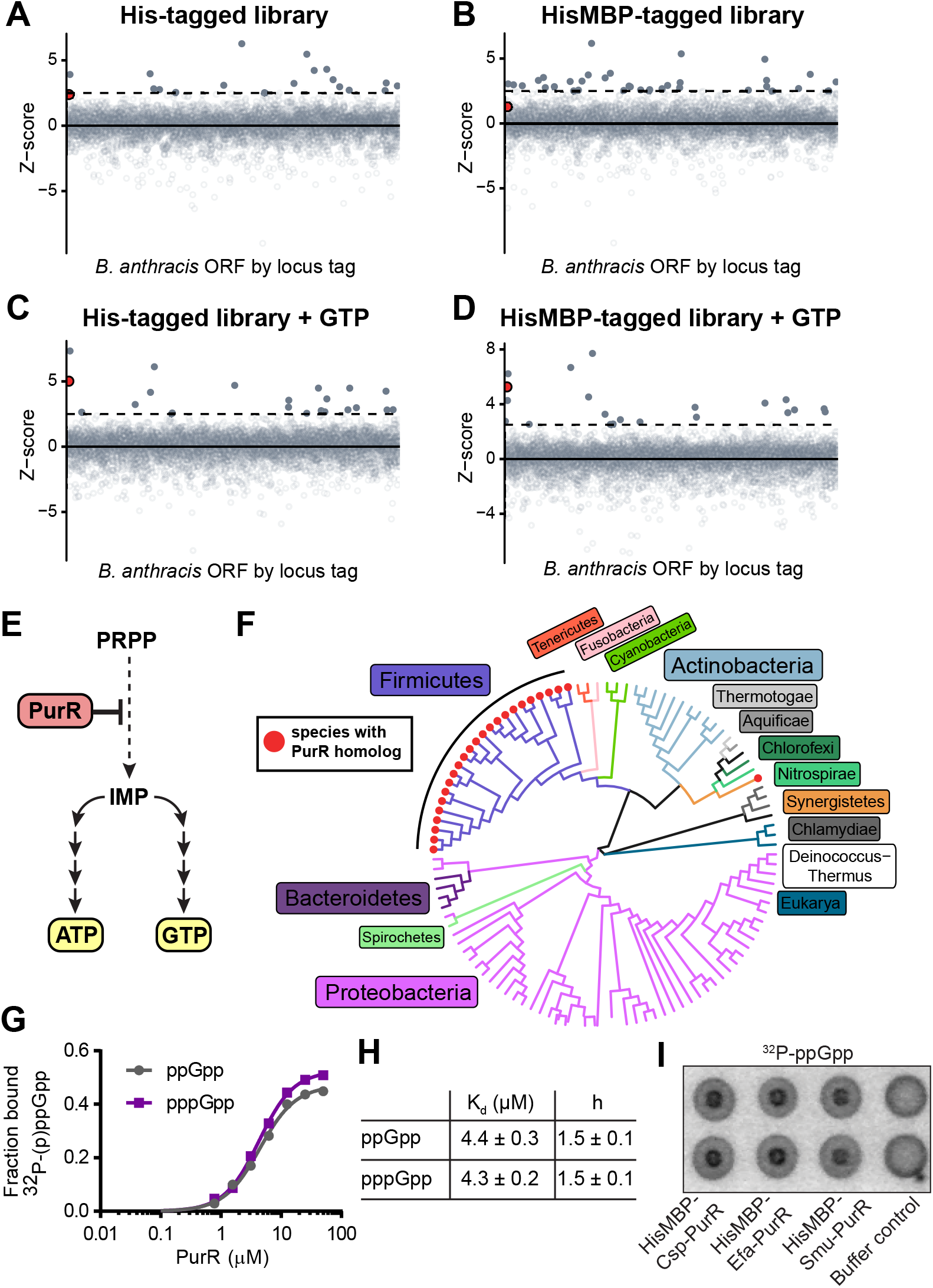
DRaCALA screen identifies the transcription factor PurR as a (p)ppGpp target. **A-B)** Z-scores of fraction ^32^P-pppGpp bound to an open reading frame (ORF) library from *B. anthracis*. His indicates hexahistidine-tagged ORFs, and HisMBP indicates hexahistidine maltose binding protein tagged ORFs. ORFs with Z-scores greater than 2.5σ are filled. Dotted horizontal line at 2.5σ. Red shows the position of the PurR ORF. **C-D)** DRaCALA of ORF libraries from (A) and (B) with 100 µM non-radioactive GTP as a competitor to reveal (p)ppGpp-specific targets. **E)** Schematic of PurR regulation of *de novo* ATP and GTP synthesis. **F)** Binding curve between purified, untagged *B. subtilis* PurR and ^32^P-ppGpp (gray) and ^32^P-pppGpp (purple) obtained with DRaCALA. Error bars, representing SEM of triplicate. **G)** Binding parameters from (E). ± standard error. h = Hill coefficient. **H)** DRaCALA of ^32^P-ppGpp binding to 30 μM purified HisMBP-tagged PurR proteins from *Clostridium sporogenes* (Csp), *Enterococcus faecalis* (Efa), and *Streptococcus mutans* (Smu). Control is protein storage buffer.

**Table 1.**
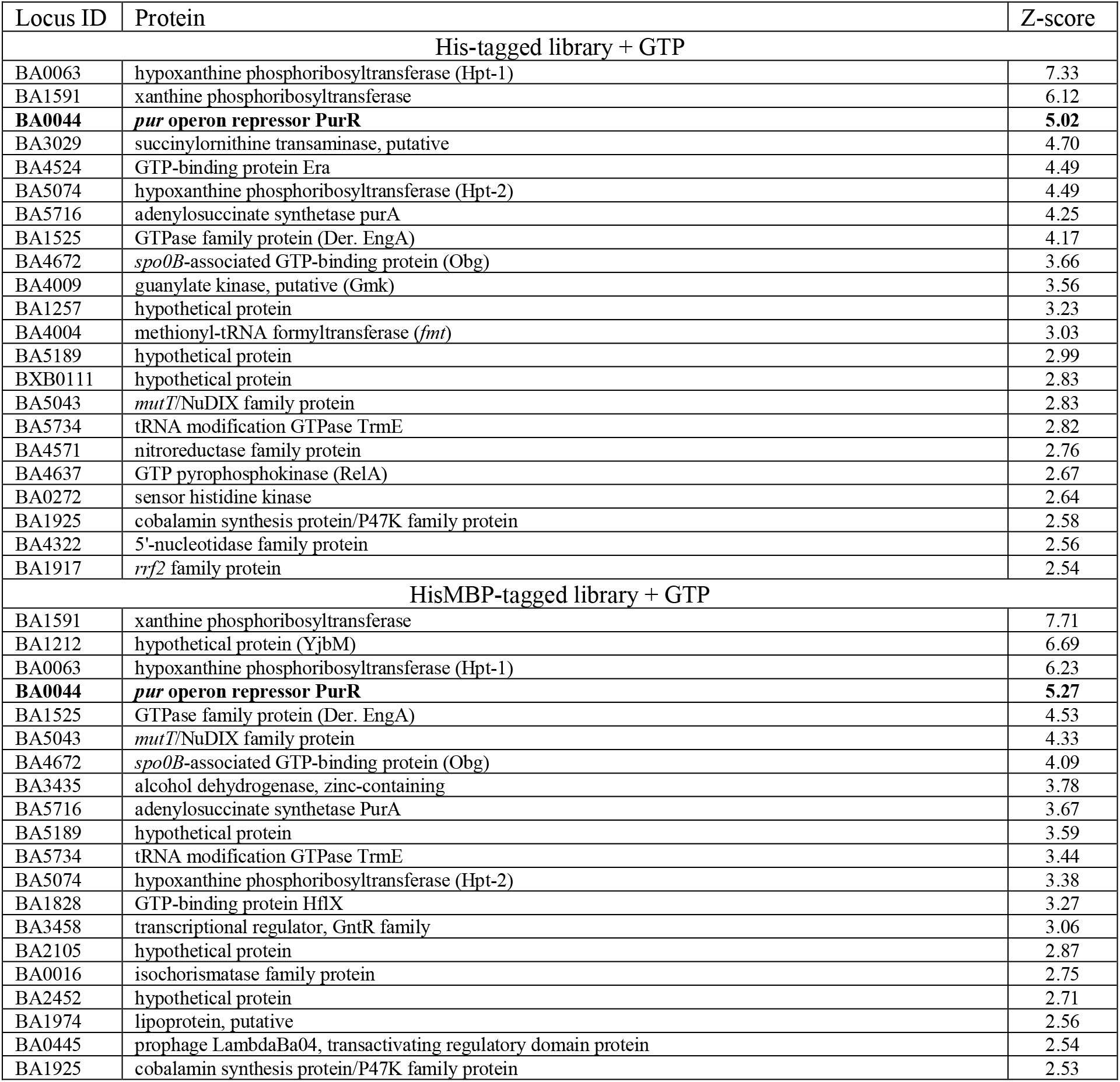
pppGpp targets identified in screening His- and HisMBP-tagged libraries of *B. anthracis* with non-radiolabeled GTP competitor.

Since (p)ppGpp binds many proteins in cell lysates, we sought to reduce nonspecific background binding by performing the same screen with a small amount (100 μM GTP) of non-radiolabeled GTP as a competitor (Figure 1C and 1D). The non-radiolabeled competitor GTP also served to enrich the screen for hits that specifically bind (p)ppGpp and not also GTP. Strikingly, the purine synthesis transcription regulator PurR (BA0044) which had a Z-score slightly below the cutoff of 2.5 in the previous screens (Figure 1A-B), emerged as one of the strongest hits in the screens with GTP, only behind the known targets HPRT, XPRT, and YjbM (SasB) (Figure 1C and 1D, Table 1, and Table S1).

PurR is a transcription regulator in bacteria, whose major role is the repression of the transcription of genes responsible for *de novo* synthesis of IMP, the common precursor to the purine nucleotides ATP and GTP (Figure 1E) (Ebbole and Zalkin, 1987; Hove-Jensen et al., 2017; Sause et al., 2019; Weng et al., 1995). PurR is a critical regulator in ensuring balanced and efficient synthesis of purine nucleotides (Rappu et al., 1999; Weng et al., 1995). Homologs of the *B. anthracis* PurR are found almost exclusively in the bacterial phylum Firmicutes (Figure 1F). PurR proteins from other bacterial phyla, such as Proteobacteria, are structurally unrelated (Choi and Zalkin, 1992; Meng and Nygaard, 1990; Schumacher et al., 1994). To further study the PurR-(p)ppGpp interaction, we turned to PurR from *Bacillus subtilis*, a close relative of *B. anthracis*, since its PurR has been extensively characterized and the PurR homologs are highly similar (64% identical) (Weng et al., 1995). We purified untagged *B. subtilis* PurR and tested for binding to ^32^P-ppGpp and ^32^P-pppGpp with DRaCALA (Figure 1G). Both ppGpp and pppGpp interacted with *B. subtilis* PurR similarly, with binding curves best fit with a Kd ∼ 4.4 μM and a Hill cooperativity coefficient of 1.5 (Figure 1G and 1H). Given that the physiological concentration of (p)ppGpp in *Bacillus* can reach up to mM, this interaction is highly relevant *in vivo*.

Given the conservation of PurR in firmicutes beyond *B. anthracis* and *B. subtilis*, we tested whether (p)ppGpp interaction extends to PurR homologs in other firmicutes, using purified PurR from the soil bacterium *Clostridium sporogenes*, the opportunistic pathogen *Enterococcus faecalis*, and the oral pathogen *Streptococcus mutans.* In all cases we detected significant binding to ^32^P-ppGpp (Figure 1I). Altogether, our screen has identified a transcription factor in Firmicutes that (p)ppGpp binds tightly and specifically. (p)ppGpp binding to PurR suggests that it can act as an effector for the transcription factor.

### (p)ppGpp binds to a conserved pocket on the effector binding domain of PurR

We next solved a co-structure of ppGpp bound to *B. subtilis* PurR to 2.45 Å resolution (Figure 2A and Table S2). *B. subtilis* PurR comprises two domains, an N-terminal winged helix-turn-helix domain for DNA binding and a C-terminal effector binding domain. In contrast to *E. coli* PurR whose effector binding domain binds purine bases and is structurally similar to a ribose binding protein, the firmicute PurR effector binding domain is similar to the phosphoribosyltransferase (PRT) class of enzymes (Figure 2A) (Schumacher et al., 1994; Sinha et al., 2003). In the structure of the *B. subtilis* PurR-ppGpp complex, ppGpp bound to the PRT domain in a pocket defined by three loops found in all PRT proteins (II, III, IV; loop numbering from PRT enzymes) (Figure 2A and 2B) (Sinha and Smith, 2001). The guanine ring of ppGpp is engaged in π-stacking interactions, sandwiched between the aromatic side chains of residues Phe205 and Tyr102 (Figure 2B). A Y102A variant interacts with ^32^P-ppGpp more weakly than wild type, supporting a role for Tyr102 in ppGpp binding (Figure S1A). The 5′-phosphates of ppGpp nestle into a pocket formed by loop III and an α-helix, where the backbone amides of loop III and the side chain of Thr211 coordinate the 5′-phosphates. The 3′-phosphates extend upward and out of the binding pocket, where they interact with backbone amides of loop II and the side chains of Tyr102 and Lys207 (Figure 2B). While (p)ppGpp is commonly co-crystallized with divalent cations (e.g. Mg^2+^ or Mn^2+^), we did not find evidence for metals in our structure. Accordingly, DRaCALA confirmed that Mg^2+^ is not required for the interaction between ^32^P-ppGpp and PurR (Figure S1B).

**Figure 2.**
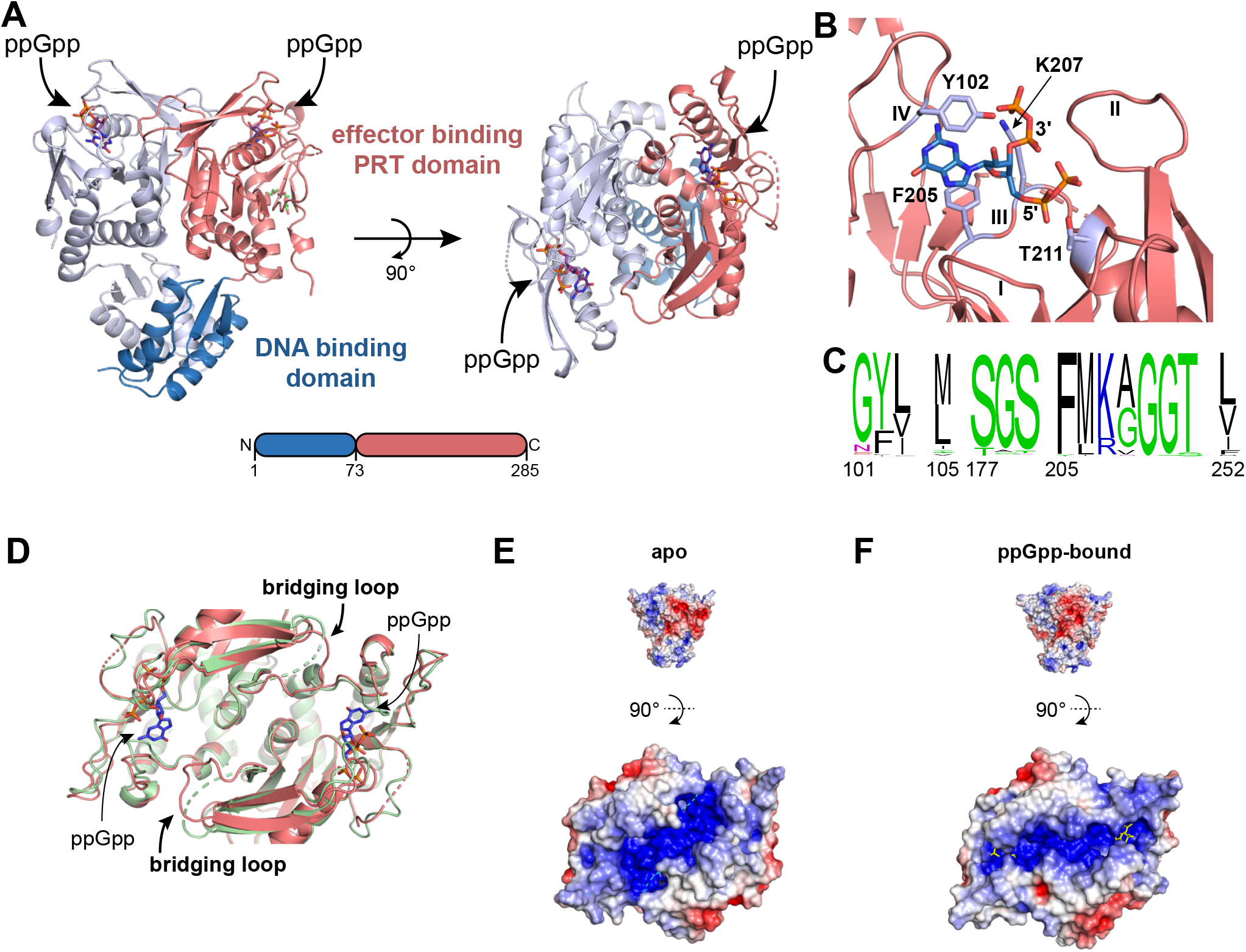
ppGpp binds to the effector binding domain of PurR. **A)** *B. subtilis* PurR dimer crystallized with ppGpp. The N-terminal DNA binding domain is blue and the C-terminal effector binding phosphoribosyltransferase (PRT) domain is salmon. (Right) Rotating 90° shows the ppGpp binding pockets on the effector domain. **B)** ppGpp binding pocket with select interacting residues indicated in silver. Loops I-IV of the PRT binding pocket are indicated. **C)** Mg^2+^ did not crystallize with PurR-ppGpp, and it is not necessary for the interaction as determined by DRaCALA. **D)** Overlay of PurR-ppGpp (salmon) with PurR-cPRPP (yellow; PDB ID 1P4A). Side chains of Y102 and D203 shown. Loop II hidden for clarity. **E)** PRPP competes with ^32^P-ppGpp binding to PurR, but PRPP competes poorly to bind PurR D203A. **F)** DRaCALA binding curves of ^32^P-ppGpp binding to Y102A and D203A variants of PurR. **G)** Overlay of apo PurR (green; PDB ID 1P4A) and ppGpp-bound PurR (salmon). Bridging loop that is unresolved in apo but not with ppGpp is indicated. **H-I)** Poisson-Boltzmann continuum electrostatics of the effector binding domain of apo (H) and ppGpp-bound (I) PurR. Scale of electrostatic potential is −5 (red) to +5 (blue).

The (p)ppGpp binding pocket among firmicutes PurR homologs is highly conserved. We aligned the (p)ppGpp binding residues from 938 *B. subtilis* PurR homologs in the Pfam database (Figure 2C). Of the 15 residues involved in the interaction, 11 are nearly perfectly conserved, and variants in other sites are highly similar (Figure 2C). For example, *B. subtilis* PurR Tyr102 is a tyrosine in most homologs and a phenylalanine in the others. Also, *B. subtilis* PurR Lys207 is a lysine in most homologs and an arginine in the remainder. Loop III residues Phe205 and Gly209/Gly210/Thr211 are nearly universally conserved. ConSurf analysis across the whole protein for the 938 PurR homologs shows that the (p)ppGpp binding pocket is more highly conserved than the remainder of the protein, even more than the DNA binding domain (Figure S1C).

Structural comparison between apo PurR and ppGpp-bound PurR does not reveal drastic conformational or electrostatic differences in the DNA binding domain (Figure S2A-C). However, ppGpp binding does affect the effector binding domain, which is important for oligomeric interaction between PurR subunits. Specifically, a loop that bridges the monomer-monomer interface of a PurR dimer is unresolved in apo PurR but is resolved in ppGpp-bound PurR, indicating that ppGpp binding stabilizes the conformation of this loop (Figure 2D). The effect of this change is that ppGpp significantly alters the electrostatics of the effector binding domain by creating a positively charged channel different from the apo protein (Figure 2E and 2F). Since it has been hypothesized that DNA wraps around PurR to interact with the effector binding domain (Bera et al., 2003), it is possible that the altered electrostatic surfaces affect the interactions with the electronegative DNA backbone.

### (p)ppGpp competes with PRPP for binding PurR’s effector binding domain to allosterically alter DNA binding

The effector binding domain of PurR is homologous to PRT enzymes, which catalyze phosphoribosyl transfer reactions with PRPP as a substrate. The PRT domain of PurR also binds PRPP but does not have enzymatic activity, instead PRPP binding de-represses the PurR regulon (Weng et al., 1995). The ppGpp binding site partially overlaps the PRPP binding site, as seen in an overlay of ppGpp and the nonreactive PRPP analog, cPRPP, bound to PurR (Figure 3A) (Bera et al., 2003). Particularly, the 5′-phosphates of ppGpp overlap with the 5′-phosphate of PRPP in their interaction with loop III (Figure 3A). The overlap in this binding site suggests that they would compete for binding to PurR. Indeed, we found PRPP competes with ^32^P-ppGpp for binding PurR (Figure 3B).

**Figure 3.**
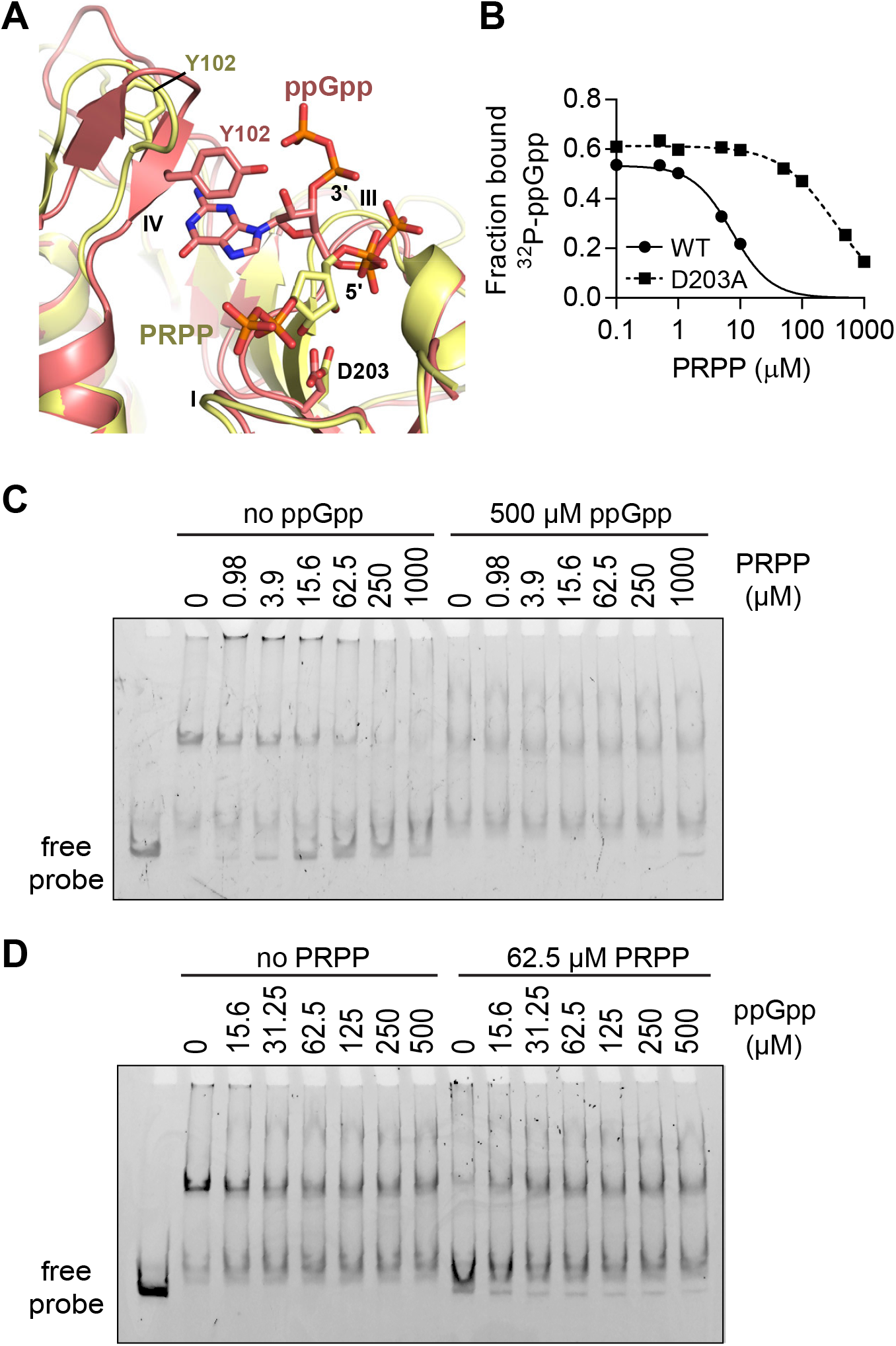
ppGpp competes with PRPP to allosterically regulate PurR-DNA interaction. **A)** Overlay of PurR-ppGpp (salmon) with PurR-cPRPP (yellow; PDB ID 1P4A). Side chains of Y102 and D203 shown. Loop II hidden for clarity. **B)** DRaCALA of competition between ^32^P-ppGpp and PRPP binding to PurR and PurR D203A. **C)** EMSA of PurR interaction with FAM-labeled 221 bp DNA with increasing PRPP concentrations and with or without ppGpp. **D)** EMSA of PurR interaction with DNA with increasing ppGpp concentration and with or without PRPP. PurR concentration is 100 nM, close to the physiological concentration of the protein. Similar EMSA results were observed with a non-labeled 202 bp probe from the same control region with a lower KCl concentration and no nonspecific DNA in the EMSA reaction (Figure S3A-D).

The binding pocket also contains interactions specific for each effector. PurR Tyr102 is flipped away from the pocket with PRPP bound and covers the guanine ring with ppGpp bound (Figure 3A). Asp203 sits below the ribose of PRPP but does not interact with ppGpp (Figure 3A). An alanine variant at this position reduced the ability of PRPP to compete with ppGpp by 100-fold (Figure 3B), but ^32^P-ppGpp binding is unaffected (Figure S1A).

To examine the role of (p)ppGpp on PurR’s ability to regulate transcription, we assessed PurR binding to its cognate promoter of the *pur* operon *in vitro* with electrophoretic mobility shift assays (EMSAs). To test the specificity of the interaction, we used a FAM-labeled 221 bp region of the promoter DNA in the presence of nonspecific DNA competitor and at high salt concentration (Figure 3C). We observed a complete probe shift induced by PurR in the absence of any ligands, indicating that PurR binds to the promoter DNA. We then performed EMSA in the presence of increasing concentrations of PRPP. As expected, the PurR-DNA interaction was noticeably inhibited by as low as <10 μM of PRPP, and completely inhibited by physiological concentration of 1000 μM PRPP, resulting in loss of the larger molecular weight complex and an increase in unbound DNA (Figure 3C). However, when ppGpp was added in competition with PRPP, it prevented PRPP from abolishing PurR-DNA interaction, and only a small portion of unbound probe was observed even at the highest PRPP concentration (Figure 3C). These results suggest that by binding the effector domain of PurR, ppGpp maintains PurR-DNA interaction and prevents PRPP from de-repressing PurR regulation.

We also performed EMSA in the presence of increasing amount of ppGpp. We noted that ppGpp alone has an effect on the PurR-DNA interaction (Figure 3D and Figure S3A-D). However, attempts to crystallize the PurR-DNA-ppGpp complex were unsuccessful. With no structural knowledge of the PurR-DNA interaction, deducing mechanistic details of ppGpp’s effect is difficult. Importantly, in the presence of even low concentration of PRPP, the addition of ppGpp greatly enhances PurR-DNA interaction (Figure 3D). Because PRPP is a key intermediate for nucleotide biosynthesis, it is always present *in vivo*. Therefore, our results suggest that (p)ppGpp is a co-repressor of PurR critical for its repression of the PurR regulon *in vivo*.

Overall, we conclude that (p)ppGpp binds PurR at an effector binding pocket distal to the DNA binding domain and competes with the inducer ligand PRPP to enhance the PurR-DNA interaction.

### Defining the PurR regulon in *Bacillus subtilis*

(p)ppGpp binding to PurR raised the possibility that (p)ppGpp controls expression of the PurR regulon during stress. Since the PurR regulon in *B. subtilis* had not been systematically characterized, we turned to ChIP-seq to fully document PurR binding sites. We raised polyclonal antibodies against untagged *B. subtilis* PurR (Figure S4) and collected ChIP samples from *B. subtilis* grown in medium containing the nucleobases adenine, guanine, cytosine, and uracil, which are known to increase PurR repression (Weng et al., 1995). ChIP-seq identified 15 PurR binding sites genome-wide, including six not previously known to be PurR binding sites in *B. subtilis* (Figure 4A-B, Figure S5 and Table 2) (Saxild et al., 2001). These PurR binding sites control expression of more than 30 genes, the majority of which are involved in purine nucleotide synthesis (Figure 4A). For example, PurR binds upstream of the 12-gene *pur* operon (*purEKBCSQLFMNHD*), which encodes all proteins necessary for *de novo* synthesis of the GTP/ATP precursor IMP from PRPP (Figure 4A and 4C). PurR also binds upstream of *pbuG* and *pbuO* (guanine/hypoxanthine permeases) and *xpt-pbuX* (xanthine phosphoribosyltransferase and xanthine permease), which altogether encode most proteins in the purine salvage pathway, and *purA* (the first dedicated step towards ATP synthesis) and *guaC* (GMP reductase that converts GMP to IMP). PurR controls expression of *nusB-folD* and *glyA*, which contribute to 10-formyltetrahydrofolate production necessary for *de novo* IMP synthesis (Figure 4B). Lastly, PurR binds upstream of its own promoter at the *purR-yabJ* operon.

**Figure 4.**
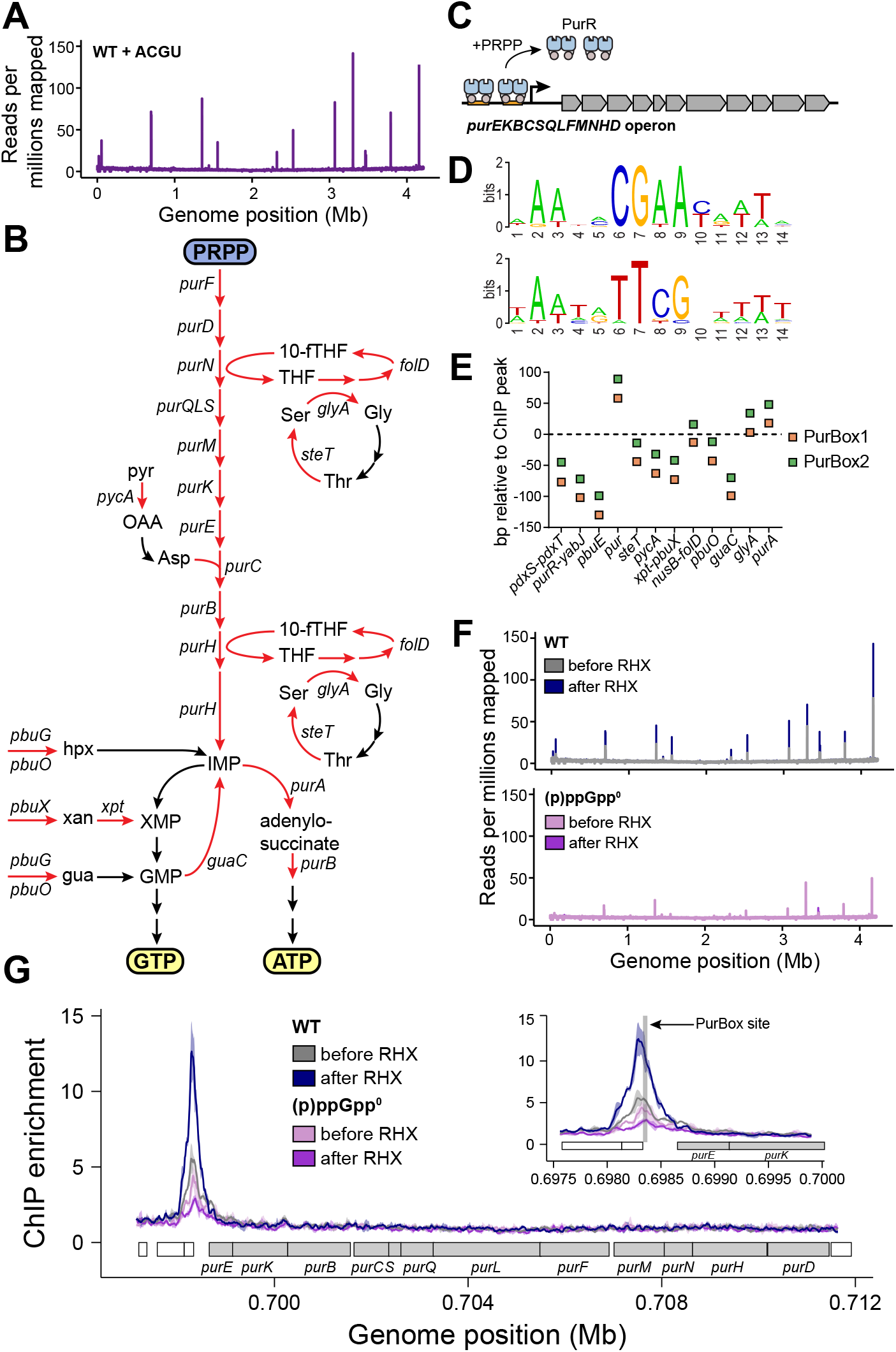
(p)ppGpp enhances PurR-DNA interaction during stress response. **A)** Genome-wide view of mean PurR ChIP reads per million reads from *B. subtilis* grown with the nucleobases ACGU. **B)** GTP and ATP synthesis pathway in *B. subtilis*. Arrows in red with gene names show genes that ChIP-seq identified in the PurR regulon. 10-fTHF=10-formyl tetrahydrofolate; ser=serine; gly=glycine; thr=threonine; pyr=pyruvate; OAA=oxaloacetate; asp=aspartate; hpx=hypoxanthine; xan=xanthine; gua=guanine. Placements of *pycA* and *steT* in this pathway are inferred based on biological function and have not been verified. **C)** Schematic of the *pur* operon and its repression by PurR in *B. subtilis*. **D)** Frequency logos of PurBox1 (top) and PurBox2 (bottom) sequences from 12 of the 15 PurR ChIP peaks. Frequency logos were created using WebLogo (UC Berkeley). **E)** Distance of PurBoxes from the ChIP enrichment peaks. Peak represented by the dotted horizontal line at Y=0. Peak location determined at 10 bp resolution from PurR ChIP sample obtained from (p)ppGpp-induced WT *B. subtilis*. Distance calculated from the center of each PurBox (7th nt out of 15 nt PurBox) to the peak location. **F)** Genome-wide view of mean PurR ChIP reads per million reads in WT and (p)ppGpp^0^. **G)** PurR ChIP enrichment at the *pur* operon in WT and (p)ppGpp^0^. Inset zooms in on the site, and shaded box shows PurR binding sequence.

**Table 2.**
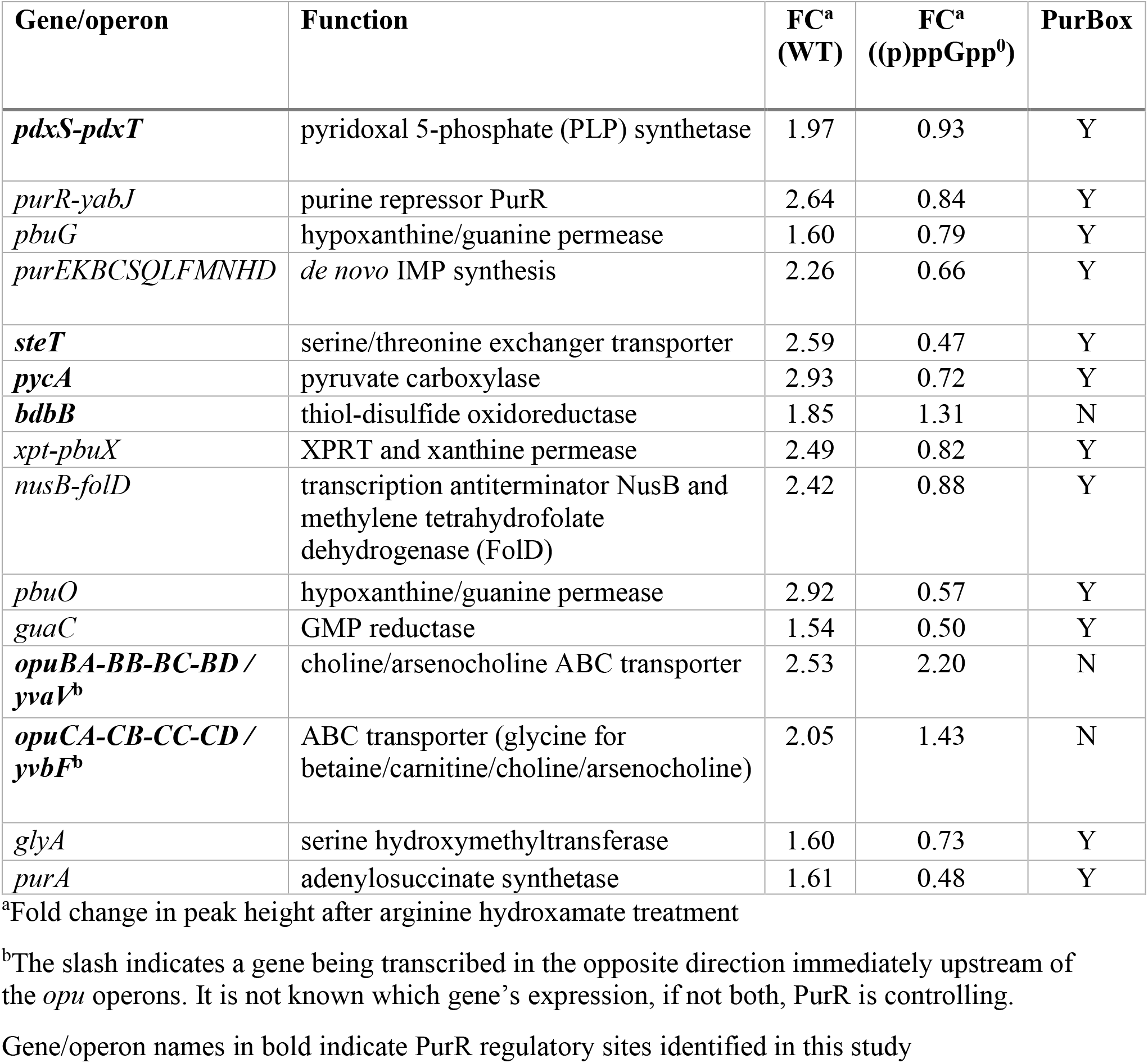
PurR binding sites in *B. subtilis* identified with ChIP-seq.

Two of the novel PurR binding sites can also be linked to purine nucleotide metabolism. The gene *steT* encodes a serine/threonine exchanger that we hypothesize may be involved in 10-formyltetrahydrofolate regeneration (Figure 4B). The gene *pycA* encodes pyruvate carboxylase, which can be linked to aspartate synthesis, which is necessary for *de novo* IMP synthesis (Figure 4B). Unexpected binding sites include upstream of the *pdxS-pdxT* operon, which encodes the enzyme complex required for *de novo* PLP (vitamin B6) synthesis (Sakai et al., 2002). There are at least 60 PLP-dependent proteins in *B. subtilis*, a majority of them involved in amino acid biosynthesis (Richts et al., 2019). Seven transcription factors bind PLP, including GabR which regulates γ-aminobutyrate utilization (Belitsky, 2004; Belitsky and Sonenshein, 2002). We also identified ChIP peaks at the *opuB* and *opuC* operons, which encode transporters involved in osmotic protection in *B. subtilis* (Hoffmann et al., 2018; Lee et al., 2013). The final PurR binding site with the weakest ChIP enrichment is intragenic upstream of *bdbB*, encoding a thiol-disulfide reductase.

PurR binding sites are characterized by two inverted repeats, termed PurBoxes, that each bind a PurR dimer (Figure 4C) (Shin et al., 1997). In addition to the PurBoxes at the nine previously known PurR binding sites (Table 2) (Saxild et al., 2001), we identified PurBoxes at three out of the six novel binding sites (Figure S5 and Table 2). A frequency logo shows that the PurBoxes at the 12 sites are characterized by a central CGAA motif (or TTCG for the inverted repeat) surrounded by A-T rich sequences (Figure 4D). Interestingly, the peak of PurR ChIP occupancy was offset from the PurBoxes (Figure 4E), suggesting that PurR protects DNA distal from the PurBox binding sequence, possibly through extended PurR oligomerization. Since PurR copy numbers range from 1,000-3,000 per cell in *B. subtilis* (Maass et al., 2011, 2014), there are ample PurR copies to extend beyond the PurBoxes at each binding site.

### (p)ppGpp is a co-repressor of the PurR regulon following starvation

Next, we examined the PurR ChIP patterns 10 minutes after amino acid starvation, which was mimicked using the drug arginine hydroxamate (RHX). RHX induces (p)ppGpp to millimolar levels in wild type cells to turn on the stringent response. Strikingly, PurR ChIP signal increased at each PurR binding site in wild type cells by an average of 2.2-fold (Figure 4F and S6 and Table 2). This included a 2.3-fold increase at the *pur* operon (Figure 4G and Table 2). On the other hand, the PurR ChIP signal remained unchanged or decreased slightly in a (p)ppGpp-null ((p)ppGpp^0^) mutant (Figure 4G and S6 and Table 2).

We isolated the effect of (p)ppGpp on the change in PurR occupancy after RHX treatment by calculating the change in global PurR occupancy after RHX in both wild type and (p)ppGpp^0^ cells and taking the ratio of the changes in wild type over (p)ppGpp^0^. We found a (p)ppGpp-dependent increase in PurR occupancy at 12 out of the 15 binding sites (Figure S7), including at the *pur* operon, indicating that (p)ppGpp is necessary for increased PurR occupancy at these sites. All sites with canonical PurBoxes had a (p)ppGpp-dependent increase in PurR occupancy, suggesting that the PurBoxes are necessary for (p)ppGpp co-repression.

In accordance with the hypothesis that PurR increases repression of the PurR regulon, we observed that many genes in the PurR regulon, including those in the *pur* operon, are downregulated up to 30-fold following amino acid starvation in *B. subtilis*, but no downregulation is observed in (p)ppGpp^0^ (Figure 5A) (Kriel et al., 2014).

**Figure 5.**
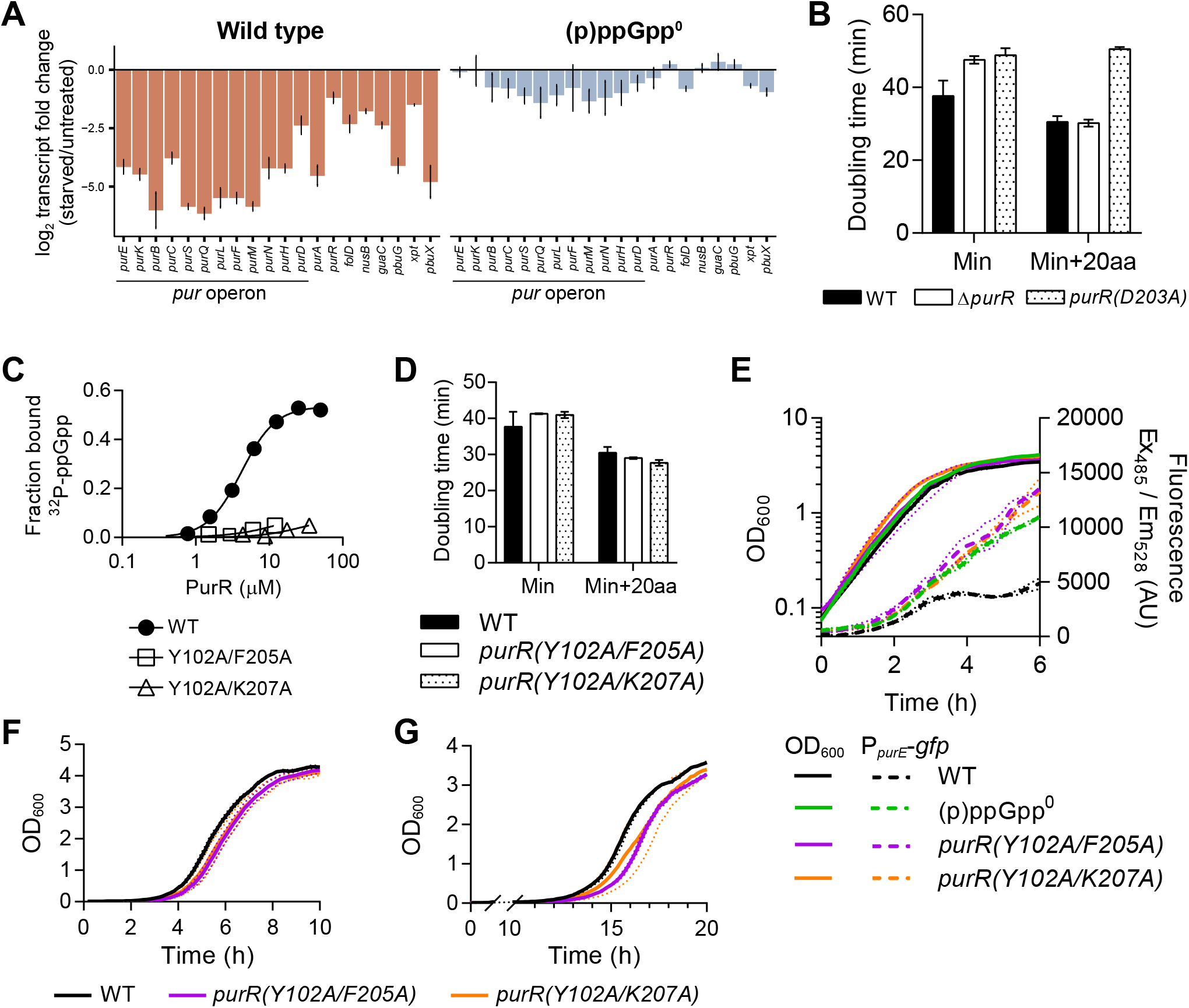
(p)ppGpp regulation of PurR is important for nutrient stress adaptation. **A)** Change in transcript level of PurR regulon after RHX treatment. Data from (Kriel et al., 2014). Mean of triplicate ± SD is shown. **B)** Doubling times of *B. subtilis* wild type, Δ*purR*, and *purR(D203A)* in minimal medium (Min) and Min supplemented with 20 amino acids (20aa). **C)** DRaCALA binding curves showing ^32^P-ppGpp interaction with PurR Y102A/F205A and Y102A/K207A. **D)** Doubling times of *purR(Y102A/F205A)* and *purR(Y102A/K207A)* in Min and Min+20aa media. **E)** Expression of a P*purE*-GFP reporter in wild type, (p)ppGpp^0^, *purR(Y102A/F205A)*, and *purR(Y102A/K207A) B. subtilis*. **F-G)** Growth of wild type, *purR(Y102A/F205A)*, and *purR(Y102A/K207A)* in Min+20aa (F) and Min (G) media following the nutrient downshift from Min+20aa to Min media. All error bars are SEM of triplicate. For OD and fluorescence curves, error bars are represented by dotted lines.

These data demonstrate that (p)ppGpp acts as a co-repressor of PurR in *B. subtilis*. By binding PurR, (p)ppGpp controls expression of 28 genes involved in purine nucleotide synthesis through both *de novo* and salvage pathways. This will lead to reduction of GTP levels, which is necessary for adaptation to nutrient and antibiotic stresses in Firmicutes.

### (p)ppGpp binding to PurR is important for nutrient stress response

Transcriptional regulation of purine nucleotide synthesis is important for controlling ATP and GTP levels. In *B. subtilis*, GTP levels correlate with growth rate, and reducing GTP is critical for surviving nutrient stress (Bittner et al., 2014; Kriel et al., 2012). We first demonstrated the importance of transcriptional regulation of purine nucleotide synthesis using a strain lacking PurR regulation (Δ*purR*) and a strain with uninducible PurR repression (*purR(D203A)* which lacks the ability to bind PRPP) (Rappu et al., 2003; Weng and Zalkin, 2000). When these strains are grown in a minimal medium requiring *de novo* nucleotide synthesis, they display increased doubling times relative to wild type *B. subtilis* (Figure 5B). Uninducible repression of purine nucleotide synthesis also caused a significant increase in doubling time in a medium replete with amino acids (Figure 5B). These results demonstrate that controlling transcription of *de novo* purine nucleotide synthesis is required for optimal growth of *B. subtilis*.

Increased repression of the *pur* operon during a stress response suggests that (p)ppGpp controls *de novo* GTP synthesis for optimal synthesis during a stress response (Figure 5A). To test the importance for (p)ppGpp binding to PurR for stress adaptation, we created two (p)ppGpp-refractory PurR variants, PurR (Y102A/F205A) and PurR (Y102A/K207A) (Figure 5C) and introduced the variants into the *B. subtilis* genome at the endogenous *purR* locus. Both variants had doubling times similar to wild type *B. subtilis* during steady state growth (Figure 5D). Notably, both variants were unable to downregulate expression of a GFP reporter controlled by the *pur* operon promoter during entry into stationary phase (Figure 5E and S8A), consistent with (p)ppGpp interaction with PurR being important for repression of the PurR regulon. We grew the (p)ppGpp-refractory *purR* mutant strains in a medium replete with amino acids and downshifted them into a medium lacking amino acids through a series of washes. We then followed outgrowth of the strains in both rich and minimal media. Both *purR(Y102A/F205A)* and *purR(Y102A/K207A)* had a lag time about 30 min longer than wild type during outgrowth in rich and minimal media following downshift (Figure 5F-G and Figure S8B-C). This increased lag is likely due to lack of PurR regulation since a similar result was observed with Δ*purR*, which is not able to repress *de novo* purine synthesis during a downshift (Figure S8D). Therefore, (p)ppGpp regulation of PurR plays a significant role for *B. subtilis* to adapt to changes in amino acid availability.

## DISCUSSION

Transcription factors can relay environmental signals to transcriptional regulation through allosteric regulation by effector ligands. Here we discovered the first example of the ubiquitous alarmone (p)ppGpp acting as an effector for a transcription repressor. Our findings are also the first demonstration of direct regulation of transcription by (p)ppGpp in Firmicutes. In firmicutes, (p)ppGpp binds the conserved transcription repressor PurR, which controls purine nucleotide biosynthesis by binding to 15 promoter sites in *B. subtilis*. (p)ppGpp competes with the PurR inducer PRPP for a binding pocket in the PurR effector binding domain, thus allosterically increasing PurR binding to the promoter DNA. This allows (p)ppGpp to serve as a co-repressor to strongly repress both *de novo* and salvage purine nucleotide synthesis during amino acid starvation and enhances bacterial survival during adaptation to fluctuating environmental conditions.

### (p)ppGpp as a classical co-repressor of a transcription regulator

Our study provides the first example of the well-characterized alarmone (p)ppGpp acting as a canonical effector of a transcription regulator. Transcription factors often bind effectors that relay external signals to internal responses, allowing a cell to modulate its physiology to adapt to environmental changes. Well-known examples include LacI binding allolactose to derepress the *lac* operon when lactose is available (Taraban et al., 2008), and TrpR binding tryptophan to enhance repression of the *trp* operon when tryptophan is already available (Zhang et al., 1987; Zhao et al., 1993). Many nucleotide second messengers also bind transcription factors to relay signals. cAMP binds CRP as a co-activator to signal low carbon availability (Sassone-Corsi, 1995). c-di-GMP binds FleQ in *P. aeruginosa* to regulate flagella expression and exopolysaccharide synthesis (Hickman and Harwood, 2008), VpsT in *Vibrio cholerae* to regulate biofilm formation (Krasteva et al., 2010), and BldD in *Streptomyces* to regulate development (Tschowri et al., 2014). c-di-AMP binds DarR in *M. tuberculosis* to influence ion and membrane homeostasis (Zhang et al., 2013). All these examples include cyclic nucleotides. However, our findings broaden regulation of transcription factors to include a linear nucleotide, the ‘magic spot’ alarmone (p)ppGpp.

Our discovery that (p)ppGpp binds PurR makes it the first case of (p)ppGpp directly regulating transcription in Firmicutes and only the third structurally-characterized example of its regulation of transcription machineries. In *E. coli*, (p)ppGpp binds the RNA polymerase at two sites, the interface of ω and β′ and the interface of β′ and the transcription factor DksA (Molodtsov et al., 2018; Ross et al., 2013, 2016). By binding these sites, (p)ppGpp is proposed to influence RNA polymerase interdomain interaction and stabilize DksA-RNA polymerase interactions (Ross et al., 2016). In *Francisella tularensis*, (p)ppGpp binds to the interface of the heterodimeric MglA-SspA complex, which interacts with the RNA polymerase to recruit the transcriptional activator PigR (Cuthbert et al., 2017). In firmicutes, (p)ppGpp does not operate by either of these mechanisms. (p)ppGpp binds PurR to enhance repression, blocking access of the RNA polymerase to the promoter region. (p)ppGpp regulates each of these transcription factors differently, revealing that there is significant diversity in the effects of (p)ppGpp on transcriptional regulation.

There is no structural information on how apo PurR interacts with DNA, but substantial indirect evidence points to DNA wrapping around a PurR dimer and PurR nucleating on the DNA. At the *pur* operon, for example, two PurBoxes each bind a PurR dimer. However, six PurR dimers can associate with this region of DNA (Bera et al., 2003; Shin et al., 1997). At over 1,000 PurR copies per cell (Maass et al., 2011, 2014), PurR far outnumbers its 15 binding sites. It is possible that additional PurR copies are nucleating on the DNA outside of the PurBoxes. Accordingly, the PurR DNase I footprint at the *pur* operon extends beyond the PurBoxes themselves, with about 60 bp upstream and 20 bp downstream protected by PurR (Ebbole and Zalkin, 1989; Weng et al., 1995). This is also confirmed in our ChIP-seq data, where the PurBoxes are offset from the highest point of ChIP enrichment at nearly all the PurR binding sites (Figure 4E). Also, PurR binding to DNA induces positive supercoiling, suggesting the PurR causes broader topological changes to DNA (Shin et al., 1997).

(p)ppGpp may have a direct effect on PurR-DNA interaction. Structural comparison suggests that (p)ppGpp binding to PurR affects its homo-oligomerization via a bridging loop (Figure 2D), which is critical for transcription factors’ function. EMSA data suggest that ppGpp-PurR associated DNA probes migrate more rapidly than apo PurR associated DNA probes (Figure 3C). Furthermore, PurR has positively charged surfaces at both the DNA binding domain and in the PRT domain near the effector binding pocket, supporting a proposal that the DNA can wrap around PurR (Sinha et al., 2003). (p)ppGpp binding the PRT domain may enhance this wrapping of the DNA and nucleation of PurR on the DNA fragment. Further structural work is needed to characterize the interaction of PurR with DNA and the effects of ppGpp and PRPP on this interaction.

It is now apparent that multiple bacterial phyla have evolved ways to use (p)ppGpp to directly regulate transcription. (p)ppGpp regulons have been recently delineated in proteobacteria. (p)ppGpp’s interaction with RNA polymerase allows it to regulate over 700 genes in *E. coli*, and its binding to MglA-SspA-PigR can upregulate at least 16 genes of the *Francisella* pathogenicity island (Charity et al., 2009; Cuthbert et al., 2017; Nano and Schmerk, 2007; Sanchez-Vazquez et al., 2019). In Firmicutes, (p)ppGpp was previously only associated with indirect changes in the expression of thousands of genes, many of them likely related to (p)ppGpp’s regulation of nucleotide enzymes and depletion of GTP (Kriel et al., 2014). These include down regulation of ribosomal RNA which is controlled by concentration of the initiating nucleotide GTP (Krásný and Gourse, 2004), and the >200 genes in the *B. subtilis* CodY regulon which uses GTP as a co-repressor. (p)ppGpp indirectly activates the CodY regulon by decreasing GTP levels (Belitsky and Sonenshein, 2013; Brinsmade et al., 2014). Here we have found that expression of 28 genes in *B. subtilis* is dependent on (p)ppGpp through its interaction with PurR (Figure S7 and Table 2). Our work shows that as nutrient deprivation signals (p)ppGpp synthesis, (p)ppGpp binds PurR as a co-repressor and represses the synthesis of purine nucleotides to conserve energy. Thus we now expand our understanding of (p)ppGpp transcriptional regulon in Firmicutes beyond indirect regulation.

### A conserved (p)ppGpp-binding motif shared among enzymes and a transcription factor

The nucleotide (p)ppGpp serves a highly pleiotropic role in protecting bacteria by binding to a multitude of protein and riboswitch targets to remodel bacterial metabolomes, transcriptomes, and proteomes and to tune bacterial physiology for growth in varied environments (Corrigan et al., 2016; Sherlock et al., 2018; Wang et al., 2018; Zhang et al., 2018). (p)ppGpp regulates its targets with a few relatively conserved direct binding motifs. In one case, (p)ppGpp binds to many GTPases through their GTP-binding motif (Buglino et al., 2002a; Fan et al., 2015; Kihira et al., 2012; Pausch et al., 2018). In another case, (p)ppGpp binds to a conserved PRPP-binding motif in the PRT fold in two enzymatic targets, the purine salvage enzymes HPRT and XPRT (Anderson et al., 2019, 2020). The (p)ppGpp binding site on PurR is the third PRT site also found to bind (p)ppGpp (Figure 6A). Here, the PRT enzyme is repurposed to become an effector binding regulatory domain of the transcription factor, the substrate PRPP is repurposed to become an inducer, and the enzyme inhibitor (p)ppGpp is repurposed to become a co-repressor (Figure 6A). This striking evolutionary strategy highlights how nature utilizes limited motifs to evolve diverse regulatory targets of (p)ppGpp, allowing it to serve as a master regulator of nutrient stress.

**Figure 6.**
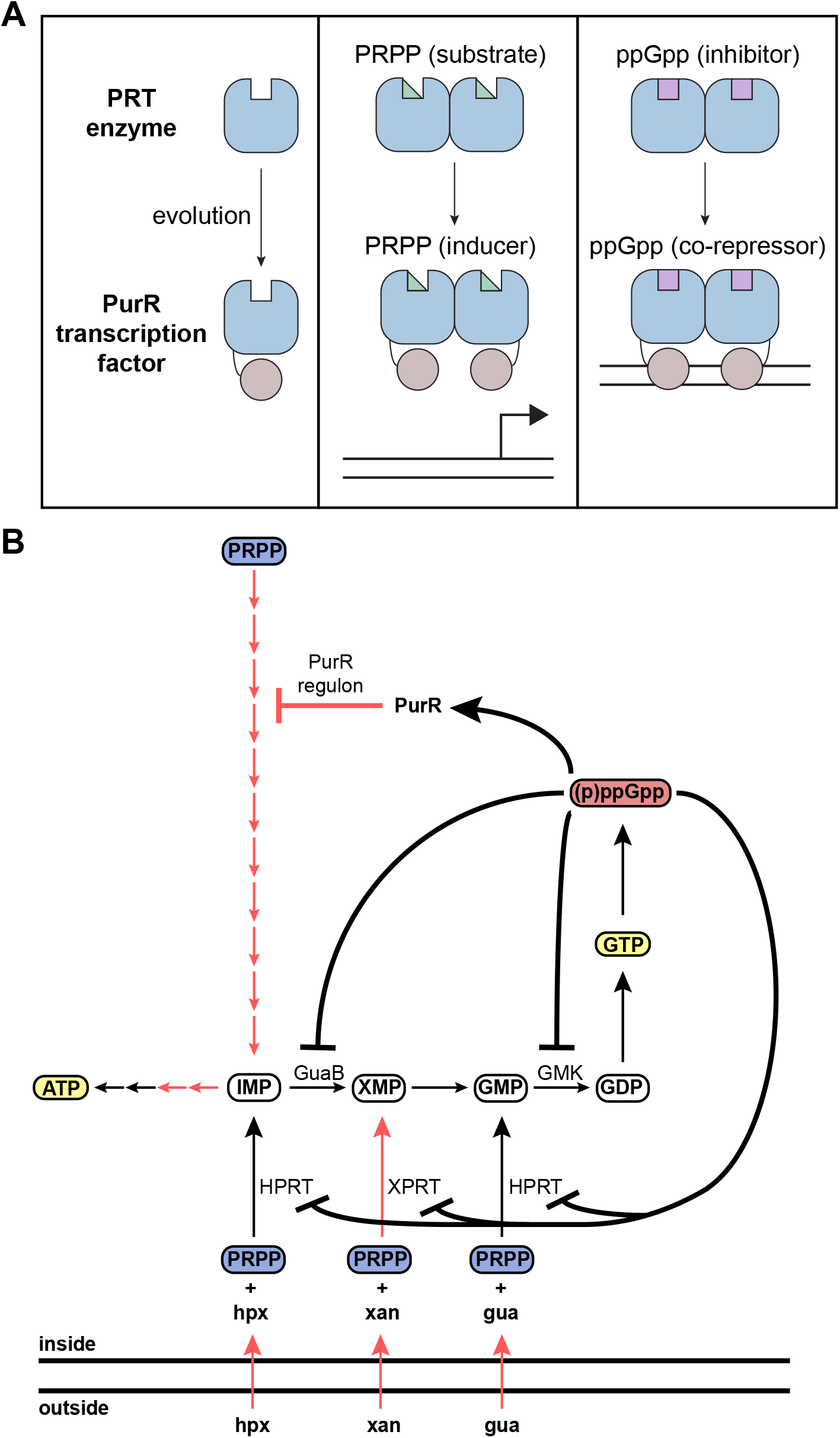
Models of (p)ppGpp regulation of PurR and global (p)ppGpp regulation of purine nucleotide synthesis in *B. subtilis*. **A)** Schematic of PurR evolution from PRT enzymes like XPRT. PurR repurposes the active site of PRT enzymes to serve as an effector binding pocket, which PRPP and ppGpp bind to allosterically regulate DNA binding. **B)** GTP and ATP synthesis in *B. subtilis*. (p)ppGpp regulates GTP synthesis at multiple points, including inhibition of GTP synthesis enzymes (GuaB, GMK, HPRT, XPRT) and binding the transcription factor PurR. Red arrows represent steps in the PurR regulon.

Despite PurR, HPRT, and XPRT sharing the same (p)ppGpp binding PRT motif, how this ligand binding site affects protein function has diversified through distinct allosteric properties. HPRT tetramerization allosterically alters the conformation of the (p)ppGpp binding pocket by holding loop II at the dimer-dimer interface (Anderson et al., 2019). In XPRT, the (p)ppGpp binding sites are cooperatively linked across a monomer-monomer interface. (p)ppGpp binding creates an electrostatic network with a bridge loop of the other monomer, inducing XPRT dimerization (Anderson et al., 2020). The PurR-ppGpp interaction introduces allosteric change of a transcription factor as a third allosteric property associated with the (p)ppGpp interacting with the same motif.

Our work has also highlighted a broader (p)ppGpp binding motif shared among GTPases, PRTs, and other targets such as guanylate kinase (Buglino et al., 2002b; DeLivron and Robinson, 2008; Kihira et al., 2012; Liu et al., 2015b). All these targets bind (p)ppGpp with a phosphate-binding loop (P-loop) motif, which is used to interact with nucleotides (Saraste et al., 1990). While the PRT proteins lack the P-loop amino acid sequence, they are structurally reminiscent of the P-loop with a loop leading into the positive dipole of an α-helix. The 5′-phosphates of (p)ppGpp and PRPP fit into the pocket created by the loop and are coordinated by the loop’s backbone (Figure 2D, 3A). This structural motif associated with (p)ppGpp binding may become more prevalent as more (p)ppGpp binding targets are characterized.

### Implication of (p)ppGpp-PRPP competition in PurR regulation

Perhaps most intriguingly, during the evolutionary co-opting of the PRT enzyme as the PurR transcription regulatory domain, not only is (p)ppGpp binding conserved, so is the (p)ppGpp-PRPP competition (Figure 6A). PRPP is the common substrates for HPRT and XPRT, and (p)ppGpp inhibits HPRT and XPRT by competing with PRPP (Anderson et al., 2019, 2020). Here we have shown that (p)ppGpp also competes with PRPP to bind PurR. In all cases, PRPP is a signal to the cell for increased nucleotide synthesis: it is directly converted to nucleotides through purine salvage or it binds PurR to de-repress the *de novo* IMP synthesis pathway (Hove-Jensen et al., 2017). (p)ppGpp competes with PRPP by binding the same binding site in each of these proteins, and (p)ppGpp shuts down nucleotide synthesis.

The competition between (p)ppGpp and PRPP for PurR binding likely tunes transcription from the PurR regulon during nutrient replete and nutrient starved growth. During unstressed growth, PRPP concentrations range from hundreds of micromolar to millimolar and (p)ppGpp is at concentrations <50 μM (Hove-Jensen et al., 2017; Kriel et al., 2012). Since both PRPP and (p)ppGpp bind PurR with Kd ∼ 5 μM (Hove-Jensen et al., 2017), in nutrient replete conditions PRPP levels dictate PurR binding and induce the PurR regulon for rapid synthesis of GTP and ATP. PRPP induction can be overridden by salvageable nucleobases, which repress the *de novo* purine operon by binding riboswitches to allow the preferential use of the salvage pathway in Firmicutes.

During amino acid starvation, (p)ppGpp levels are elevated up to >1 mM and (p)ppGpp should outcompete PRPP to strongly repress purine nucleotide synthesis. This not only conserves energy but also antagonizes the production of GTP to promote growth stasis and survival. (p)ppGpp regulation of purine nucleotide synthesis is likely critical for the coordinated use of PRPP, which is a common precursor of nucleotides (GTP, ATP, UTP) and several amino acids (Hove-Jensen et al., 2017). In *E. coli*, a mutant for which (p)ppGpp fails to inhibit purine nucleotide synthesis results in PRPP depletion due to over-production of purine nucleotides (Wang et al., 2020). (p)ppGpp inhibition of purine synthesis redirects PRPP from purine nucleotide synthesis to UTP and amino acid synthesis during starvation (Petchiappan and Gottesman, 2020; Wang et al., 2020). We propose that, in firmicutes, the direct competition between (p)ppGpp and PRPP for PurR, HPRT and XPRT (see Figure 6B) would serve the same purpose, allowing (p)ppGpp to coordinate the consumption of PRPP between purine synthesis and amino acid synthesis during starvation. Interestingly, the (p)ppGpp-PRPP competition in *B. subtilis* would allow maintenance of PRPP levels: when (p)ppGpp accumulates after amino acid starvation, it directly prevents PRPP depletion by inhibiting its consumption in purine salvage and by preventing PRPP from activating *de novo* IMP synthesis.

### (p)ppGpp controls *de novo* purine synthesis by targeting different protein factors between bacterial species

Our results reveal that (p)ppGpp regulates *de novo* purine synthesis across bacteria. The 11 reactions that synthesize IMP from PRPP are necessary for *de novo* synthesis of ATP and GTP. Until recently, there was no evidence that (p)ppGpp directly regulated this pathway in bacteria. In *E. coli*, it was recently shown that (p)ppGpp inhibits the activity of PurF, the first step in the pathway, and (p)ppGpp binding the RNA polymerase decreases the transcription of all the genes in the pathway (Sanchez-Vazquez et al., 2019; Wang et al., 2018). We have shown that (p)ppGpp regulation of *de novo* purine synthesis extends to Firmicutes, albeit by an entirely different mechanism. While Proteobacteria and Firmicutes have evolved different mechanisms for this regulation, they share the common theme of decreasing *de novo* synthesis upon (p)ppGpp induction. This theme is likely conserved across other bacteria as well.

With our data, we can begin building a model of (p)ppGpp as a global regulator of purine nucleotide synthesis in Firmicutes. As we have shown, it regulates *de novo* and salvage/scavenge nucleotide synthesis genes via its interaction with PurR (Figure 6B). (p)ppGpp also inhibits the activities of nucleobase salvage enzymes HPRT and XPRT (Anderson et al., 2019, 2020; Kriel et al., 2012), and it potently inhibits guanylate kinase activity, which catalyzes GMP to GDP (Figure 6B) (Liu et al., 2015b). Altogether, purine nucleotide synthesis has evolved to be sensitive to (p)ppGpp regulation at almost every step. Since purine nucleotides are often a signal to increase growth, (p)ppGpp counteracts this during stressful conditions to control growth and enable survival.

## MATERIALS AND METHODS

### Growth conditions

*B. subtilis* strains were grown at 37 °C unless otherwise noted. Media used to grow *B. subtilis* included lysogeny broth (LB) and S7 defined minimal medium with 1% glucose (Vasantha and Freese, 1980). S7 was supplemented with 20 amino acids (Kriel et al., 2014) and 1X ACGU (Teknova) as needed. For experiments, a single colony of each strain was resuspended in 1X Spizizen salts (Spizizen, 1958) and spread on modified Spizizen minimal agar plates (1X Spizizen salts, 1% glucose, 2 mM MgCl2, 0.7 mM CaCl2, 50 mM MnCl2, 5 mM FeCl3, 1 mM ZnCl2, 2 mM thiamine, 1.5% agar, and 0.1% glutamate). The minimal agar plates were supplemented with specific amino acids or 0.5% casamino acids where needed, especially in the case of amino acid auxotrophic (p)ppGpp^0^. The strains were grown until small colonies formed (≈16 hrs at 25-30 °C). Cultures were collected from the overnight plates with appropriate medium for the experiment and diluted to an appropriate OD600 for growth in an experiment.

Growth curves were performed with a Synergy 2 plate reader (BioTek) to measure OD600 and GFP fluorescence (Ex485/Em528). To measure growth rate, cells were washed from overnight plates, resuspended in S7 medium with 20aa at OD600 0.005, grown to OD600 ∼ 0.2, and diluted to OD600 0.005 in a clear 96-well plate (Corning) for measuring OD600 in the plate reader. Doubling times were calculated using a custom Python script that measured the fastest growth rate in a one hour window. To measure GFP fluorescence from the P*purE*-sfGFP reporter, S7 medium with 20aa in a clear bottom black 96-well plate (Corning) was inoculated with part of a single colony of bacteria. Curves of OD600 and fluorescence were translated to a starting OD600 0.075 for comparison across strains. To perform nutrient downshifts, cells were washed from overnight plates, resuspended in S7 medium with 20aa at OD600 0.01, and grown to OD600 ∼ 0.2. The medium was then filtered out with Costar Spin-X centrifuge filters (Corning, 0.45 μm) by centrifuging at 5,000 x*g* for one min. The cells were washed three times with S7 medium without any amino acids (except for glutamate), resuspended in the same medium, and diluted in a 96-well plate at OD600 0.005 in both S7 medium with and without 20aa.

### ORFeome DRaCALA screen

*Bacillus anthracis* Gateway® Clone Set containing plasmids bearing *B. anthracis* open reading frames was acquired from BEI Resources and used for Gateway cloning (Invitrogen protocol) into overexpression vectors pVL791 (10xHis tag, ampicillin-resistant) and pVL847 (10xHis-MBP tag, gentamycin-resistant) and transformed into *Escherichia coli* BL21 lacI^q^ to produce two open reading frame proteome over-expression libraries (ORFeome library).

Cell lysate with overexpressed protein and purified protein were used for DRaCALA as described (Roelofs et al., 2011). 10 μL cell lysate or diluted, purified protein was mixed with 10 μL diluted [5′-α-^32^P]-pppGpp (∼0.2 nM) in a buffer containing 10 mM Tris pH 7.5, 100 mM NaCl and 5 mM MgCl2, incubated at room temperature for 10 min. ∼2 μL mixture was blotted onto nitrocellulose membrane (Amersham; GE Healthcare) and allowed for diffusion and drying. The nitrocellulose membrane loaded with mixture was exposed on phosphor screen, which was scanned by a Typhoon FLA9000 scanner (GE Healthcare). Fraction of (p)ppGpp binding was analyzed as described (Roelofs et al., 2011).

### DRaCALA

[5′-α-^32^P] pppGpp and ppGpp were purified as previously described (Anderson et al., 2019, 2020). For PurR DRaCALA, protein in storage buffer was diluted to the appropriate concentration in 20 mM HEPES pH 8, 100 mM NaCl. ^32^P-labeled pppGpp or ppGpp was added in a final dilution of ∼1:50 from its stock. The protein and ligand were incubated for 10 min and spotted onto Protran BA85 nitrocellulose (0.45 μm, GE Healthcare) with a replicator tool (VP 408FP6S2; V&P Scientific, Inc.). After the spots dried, the nitrocellulose was exposed to a phosphor screen for at least three hours and scanned with a Typhoon FLA9000 phosphorimager. Images were analyzed with ImageJ and fraction bound ^32^P-labeled (p)ppGpp was calculated as described (Roelofs et al., 2011). Data were analyzed with GraphPad Prism v7, and binding curves were obtained by fitting the data to the equation Y = (Bmax × X^h^) / (Kd^h^ + X^h^), where h represents the Hill cooperativity coefficient.

### Protein purification

PurR proteins were expressed in *E. coli* BL21 T7 Express *lacI^q^* (New England Biolabs) from the pLIC-trPC-HMA plasmid with an N-terminal hexahistidine-maltose binding protein (HisMBP) tag. Seed cultures at OD ∼ 0.5 were diluted 1:50 into Terrific broth and grown to OD ∼ 1.5 at 37 °C prior to induction with 1 mM IPTG for four hours. Cultures were centrifuged at 10,000 x*g* for 30 min, the pellet was washed with 25 mL 1X phosphate-buffered saline (PBS), and the pellet was stored at −80 °C.

Pellets were resuspended in Lysis Buffer (25 mM Tris-HCl pH 7.5, 300 mM NaCl, 10 mM imidazole; ∼20 mL Lysis Buffer per 1 L cell pellet) with Pierce protease inhibitor tablets (ThermoFisher) and DNase I (MilliporeSigma; 500 U/L cell pellet). Cells were lysed via French press and centrifuged at 30,000 x*g* for 30 min. The supernatant was filtered through a 0.45 μm filter. HisMBP-tagged PurR was loaded onto a HisTrap FF column (GE Healthcare) equilibrated with Lysis Buffer on an AktaPure FPLC (GE Healthcare). The column was washed with 15 column volumes (CV) of Lysis Buffer with 5% Elution Buffer (25 mM Tris-HCl pH 7.5, 300 mM NaCl, 500 mM imidazole). Protein was eluted in a gradient elution from 5-50% Elution Buffer. HisMBP-tagged PurR was dialyzed with tobacco etch virus (TEV) protease overnight in 25 mM Tris-HCl pH 8, 200 mM NaCl, 1 mM dithiothreitol (DTT), 0.5 mM ethylenediaminetetraacetic acid (EDTA), and 10 mM (NH4)2SO4. Following cleavage of the HisMBP tag with TEV protease, dialyzed protein was transferred into Lysis Buffer with a HiPrep 26/10 desalting column (GE Healthcare) and passed over a HisTrap FF column. Flowthrough was collected as untagged PurR. Untagged PurR was further purified via HiPrep 26/60 Sephacryl S-200 gel filtration (GE Healthcare) equilibrated in 20 mM HEPES pH 8, 300 mM NaCl, and 50 mM (NH4)2SO4. The peak corresponding to PurR was collected, concentrated, and concentration was measured with the Bradford assay. Typical concentrations were 10-20 mg/mL and yields were ∼3-4 mg PurR/L culture. Protein was flash-frozen in liquid nitrogen and stored at −80 °C.

For small scale purification of HisMBP-tagged PurR, protein was expressed the same as above in a 5 mL volume. The cell pellet was stored at −20 °C until used. Cells were resuspended in one mL lysis buffer (20 mM sodium phosphate pH 8, 500 mM NaCl, and 10 mM imidazole). Cells were incubated with 1.6 mg lysozyme (MilliporeSigma) and 360 U Benzonase endonuclease (MilliporeSigma) for about one hour on ice for lysis. Lysate was centrifuged at 20,000 x*g* for 15 min. Supernatant was incubated with His Mag Sepharose beads (equilibrated with lysis buffer according to manufacturer’s instructions; GE Healthcare) for 30 min at 4 °C rotating end-over-end. Beads were washed three times with 500 μL wash buffer (same as lysis buffer but with 40 mM imidazole). Protein was eluted three times with 250 μL elution buffer (same as lysis buffer but with 500 mM imidazole). Eluted protein was dialyzed into 20 mM HEPES pH 8, 300 mM NaCl, 20 mM (NH4)2SO4, concentration was measured with A280 and extinction coefficient, and flash frozen with liquid nitrogen for storage at −80 °C.

### X-ray crystallography

To obtain PurR-ppGpp crystals, the purified PurR protein was concentrated to 20 mg/mL and ppGpp was added to a final concentration of 1 mM. Crystal screens were performed using the hanging drop vapor diffusion method. The best crystals were obtained by mixing this complex 1:1 with a crystallization reservoir consisting of 0.1 M 2-(*N*-morpholino)ethanesulfonic acid (MES) pH 6.5 and 18% PEG 1500. The crystals grew at room temperature and took from 5 days to 1 week to reach optimal size. The crystals take the triclinic space group P1 and diffracted beyond 2.5 Å on synchrotron sources. The crystals were cryo-protected by dipping the crystals in a 1 μL drop consisting of the crystallization reagent supplemented with 15% glycerol for 2s. The crystal was then placed directly in the cryo-stream. X-ray intensity data were collected at the Advanced Light Source (ALS) beamline 8.3.1. The data were integrated in MOSFLM and scaled using SCALA (Leslie, 2006). Phaser in CCP4 was used to solve the structure by Molecular replacement using a dimer of the 1P4A *B. subtilis* PurR structure with the ligand removed. Phaser readily located the 3 PurR dimers in the crystallographic asymmetric unit. Phenix (Adams et al., 2010) was then employed to perform refinement. After several rounds of refinement and refitting and optimizing of the model, density was evident for ppGpp molecules in 4 of the 6 PurR subunits; there was only weak density for the ppGpp molecules in the other 2 subunits. After fitting the ppGpp molecules, refinement in Phenix commenced and in the final stages of refinement ordered solvent molecules were added. The final model has Rwork/Rfree values of 19.4%/23.9% to 2.45 Å resolution and validation in Molprobity placed the structure in the top 98% of structures solved to similar resolutions.

### Electrophoretic mobility shift assay (EMSA)

Gel shift assays were performed with untagged *B. subtilis* PurR, ppGpp, PRPP, and a 221 bp FAM-labeled DNA probe or a 202 bp unlabeled DNA probe. The 221 bp probe was amplified from upstream of the *pur* operon with oJW4028/4029, and the 202 bp probe was amplified with oJW1285/1286. DNA concentration was measured with Quant-IT (Thermo Scientific). Protein was diluted from frozen stock to appropriate concentrations in 10 mM HEPES pH 8.0, 100 mM KCl, and 10% glycerol. For reactions with the FAM-labeled probe, the final reaction buffer was 10 mM HEPES pH 7.5, 200 mM KCl, 10% glycerol, 0.2 mM EDTA, 5 mM MgCl2, 1 mM DTT, 0.5 mg/mL bovine serum albumin, and 100 nM salmon sperm DNA (Invitrogen). The final DNA concentration was 5 nM. For reactions with the unlabeled 202 bp probe, the conditions were the same except 50 mM KCl, no nonspecific DNA, and one nM DNA probe. Twenty microliter reactions were prepared from a 5X buffer, 10X protein stock, 10X DNA probe stock, and 10X ppGpp or PRPP stock. Reactions were mixed by pipetting and incubated at room temperature (∼23 °C) for 30 min. A 6% TBE polyacrylamide gel (Novex, Thermo Scientific) was pre-run at 10 V for 10 min at 4 °C. Ten microliters of the sample were loaded and the gel was run at 100 V for ∼100 min at 4 °C. For the unlabeled 202 bp probe, gels were stained with 1X SYBR Gold (Thermo Scientific) at room temperature for 30 min and imaged on a Typhoon FLA9000. For the FAM-labeled probe, gels were imaged on a Typhoon with FAM detection settings.

### ChIP-Seq

The PurR ChIP protocol was adapted from previously published ChIP protocols (Bonocora and Wade, 2015; Zhang et al., 2014). For sample collection, cells were grown in S7 minimal medium supplemented with the amino acids VILMTHE from a starting OD ∼ 0.01. Culture volume was chosen based on 50 mL culture being needed for each sample and timepoint collected. Cells were grown to an OD ∼ 0.5 at 37 °C and 250 RPM. An untreated sample was collected by adding ∼50 mL culture into 1.4 mL 37% formaldehyde in a 50-mL conical. The conical was shaken at 37 °C for 7-8 minutes prior to placing on ice. The remaining culture was treated with 0.5 mg/mL arginine hydroxamate (RHX) and another sample was crosslinked 10 min after RHX addition. Crosslinked samples were held on ice for at least 30 min, centrifuged at 5,000 x*g* for 15 min, washed twice with ice-cold 25 mL 1X PBS, and resuspended in 1 mL ice-cold Solution A (20% sucrose, 10 mM Tris pH 8, 10 mM EDTA, 50 mM NaCl). Samples were split into two equal volume microcentrifuge tubes and stored at −80 °C.

For chromatin immunoprecipitation, one tube of each sample was thawed and treated with 1 mg/mL lysozyme (MilliporeSigma) at 37 °C for 30 min. Lysed sample was diluted with 600 μL 2X IP buffer (100 mM Tris pH 7, 300 mM NaCl, 10 mM EDTA, 2% Triton X-100) and 0.5 mM AEBSF and divided into two tubes ∼550 μL each. Samples were sonicated with a Misonix S-4000 at amplitude 80 for 10 s ON, 15 s OFF, and a total ON time of 8-10 min. Like samples were recombined and 100 μL of sonicated sample was removed as the input sample. To each sonicated sample, 10 μL of polyclonal α-*B. subtilis* PurR serum (serum from terminal bleed of rabbit WI606 raised by Covance) was added. Antibody and lysate were incubated end-over-end overnight at 4 °C. Antibody/protein mixture was added to 50 μL Dynabeads Protein A (ThermoFisher) and incubated end-over-end for one hour at 4 °C. Beads were washed four times with lysis buffer 150 (50 mM HEPES-KOH pH 7.5, 150 mM NaCl, 1 mM EDTA, 1% Triton X-100, 0.1% SDS, 0.1% sodium deoxycholate) and two times with 10 mM Tris pH 7.5. See (Bonocora and Wade, 2015) for details on washes. Beads were resuspended in a 100 μL blunt enzyme mix reaction (Quick Blunting Kit, NEB) with half the enzyme concentration of the manufacturer’s protocol. The blunting and 5′ phosphorylation reaction occurred at 24 °C for 30 min with gentle rotation. Beads were then washed twice with lysis buffer 150 and twice with 10 mM Tris pH 8. A-tails were added to the 3′ ends of the DNA by incubating the beads with Klenow fragment (exo-) (NEB) in a 100 μL reaction. Following 30 min incubation at 37 °C, beads were washed twice with lysis buffer 150 and twice with 10 mM Tris pH 7.5. Annealed adapters were ligated onto DNA by resuspending the beads in a 100 μL reaction containing Quick Ligase (NEB) and 0.1 μM annealed i5/i7 adapter. The ligation reaction was incubated for 15 min at 24 °C. The beads were washed twice with lysis buffer 150, once with lysis buffer 500 (same composition as lysis buffer 150 except 500 mM NaCl), once with ChIP wash buffer (10 mM Tris pH 8, 250 mM LiCl, 1 mM EDTA, 0.5% sodium deoxycholate, 0.5% Nonidet-P40 (substitute) (Dot Scientific)), and once with TE buffer. Beads were resuspended in 48 μL elution buffer (50 mM Tris pH 8, 1 mM EDTA, 1% SDS) and 1.6 U Proteinase K (NEB), incubated at 55 °C for one hour and then 65 °C overnight. Supernatant was transferred to a new tube and an additional 19 μL elution buffer and 0.8 U Proteinase K were added to the beads. Following incubation at 55 °C for one hour, reverse-crosslinked samples were combined and the DNA was purified using solid phase reverse immobilization (SPRI) Sera-Mag SpeedBeads (1.6:1 ratio, GE Healthcare). The final library was eluted with 22 μL ddH_2_O.

Input samples were processed by first adding 0.5 mg/mL RNase A (ThermoFisher) and incubating at 37 °C for at least one hour. Then, 48 μL elution buffer and 1.6 U Proteinase K were added, incubated at 55 °C for one hour, and then at 65 °C overnight to reverse crosslinks. The DNA was isolated with a 1.8:1 SPRI cleanup and the concentration was quantified with Quant-iT (ThermoFisher). Moving forward, 100 ng input DNA was processed with reactions similar to the ChIP samples, except the reaction volumes were 50 μL. Following blunting/5′ phosphorylation and A-tailing, the DNA was purified with a 1.8:1 SPRI cleanup. Following ligation, the DNA was purified with two successive 1:1 or 1.2:1 SPRI cleanups to eliminate adapter carryover. The final library was eluted with 22 μL ddH_2_O.

Quantitative PCR was used to determine the number of cycles for library amplification. In a 20 μL reaction, 2 μL of each sample DNA was combined with 0.2 μM each of oJW3536 and oJW3537, 1X EvaGreen dye (Biotium), and Kapa high-fidelity low-bias polymerase (Kapa). Samples were denatured in Bio-Rad CFX qPCR thermocycler at 95 °C for 2 min followed by 40 cycles of 95 °C for 10 s, 55 °C for 10 s, and 72 °C for 30 s. The cycle quantification (Cq) value provided by the program was used as the number of cycles for full library amplification. The remaining 20 μL of library was amplified with Kapa polymerase and the primers and concentrations listed above. The reactions were denatured at 98 °C for 30 s, cycled the Cq value through 98 °C for 10 s, 55 °C for 15 s, and 72 °C for 30 s before a final extension of 72 °C for 5 min. Final amplified libraries were purified with a 1.6:1 SPRI cleanup and eluted with ddH_2_O.

### Conservation of PurR and the (p)ppGpp binding pocket

Conservation analysis of the protein and binding pocket was conducted on 938 PurR homologs in the Pfam family PF09182. Amino acid sequences were downloaded from UniProt, and proteins were aligned with MUSCLE in MEGA X. A frequency logo was created from aligned (p)ppGpp binding residues using WebLogo (UC Berkeley). ConSurf analysis was conducted using the ConSurf server (http://consurf.tau.ac.il/) with conservation score determined by Bayesian inference (Ashkenazy et al., 2016; Landau et al., 2005). Conservation scores were mapped onto PDB ID 1O57.

## SUPPLEMENTAL MATERIALS AND METHODS

### Cloning and strain construction

Primers, plasmids, and strains used in this study are listed in Table S3, Table S4, and Table S5. For selection in *B. subtilis*, media was supplemented with antibiotics when required: MLS (erythromycin at 0.5 μg/mL and lincomycin at 12.5 μg/mL) and spectinomycin (80 μg/mL). For *E. coli*, carbenicillin was used at 100 μg/mL.

*B. subtilis purR* was cloned into pLIC-trPC-HMA. PurR variants were made by megaprimer site-directed mutagenesis of the plasmid (Kirsch and Joly, 1998). Inserts and mutants were confirmed by PCR amplification and Sanger sequencing.

*B. subtilis* NCIB 3610 *purR::ermHI* was constructed by amplifying the *purR::ermHI* genomic region from *B. subtilis* 168 *purR::ermHI* (BKE00470, *Bacillus* Genetic Stock Center). The PCR product was transformed into JDW2809, and transformants were selected for MLS resistance. Positive transformants were confirmed by PCR.

To construct *purR(D203A)* and *purR(Y102A)* mutants in *B. subtilis*, CRISPR-Cas9 plasmids were cloned with Golden Gate assembly of pJW557 (amplified with primers oJW2775/2821), the appropriate repair template (gene fragment from GeneWiz), and the appropriate guide RNA (annealed oligos from IDT). Plasmids were transformed into JDW159 (*recA^+^ E. coli*) with selection on carbenicillin. Purified plasmid was transformed into JDW2809 and transformants were selected for spectinomycin resistance. Transformants were re-streaked twice on LB at 45 °C to cure the plasmid. Transformants were patched on spectinomycin, and the *purR* region of spectinomycin-sensitive colonies was sequenced to confirm the mutation. To construct the *purR(Y102A/F205A)* and *purR(Y102A/K207A)* double mutants, CRISPR-Cas9 plasmids were constructed with repair templates for *purR(F205A)* and *purR(K207A*) with the same guide RNA as *purR(D203A)*. These plasmids were transformed into *purR(Y102A)*.

To construct P*purE*-sfGFP reporters, 500 bp DNA upstream of the start codon of *purE* in *B. subtilis* was amplified with primers oJW3438/3439 and cloned into pJW669 with Golden Gate assembly according to New England Biolabs recommended protocol. The plasmid was transformed into JDW159 (*recA^+^ E. coli*) with selection on carbenicillin. Insert was confirmed by sequencing. Purified plasmid was transformed into JDW2809, JDW3970, JDW3975, and *B. subtilis* Δ*yjbM* Δ*ywaC*. Transformants were selected on spectinomycin and streaked twice on LB at 45 °C to cure the plasmid. Spectinomycin-sensitive colonies were confirmed by PCR. (p)ppGpp^0^ with P*purE*-sfGFP was constructed by a transforming *relA*::*mls* PCR fragment amplified with primers oJW902/903 into *B. subtilis* Δ*yjbM* Δ*ywaC amyE*::P*purE*-sfGFP (Kriel et al., 2012). Transformants were selected for MLS resistance. Transformants were confirmed by PCR.

P*ilvB*-GFPmut2^A206K^ reporters were constructed as previously described (Fung et al., 2020). Briefly, linearized pJW558 was transformed into JDW2809, JDW3970, and JDW3975 and selected for spectinomycin resistance. Transformants were confirmed with PCR.

### Western blot analysis

To test for α-PurR antibody activity, cultures were obtained by inoculating LB with single colonies of wild type (JDW2809) and *purR::ermHI B. subtilis* (JDW3359). Ten mL of culture were pelleted at 4500 x*g* for 20 min and washed with one mL of 50 mM Tris pH 7.5 and 300 mM NaCl. The pellet was resuspended in 100 μL of lysis buffer (10 mM Tris pH 7.5, 10 mM EDTA, 0.1 mg/mL lysozyme, 0.5 mM PMSF). Cells were lysed by incubation at 37 °C for 30 min. An equal volume of 2X Laemmli buffer (Bio-Rad) was added and cells were incubated at 95 °C for 10 min. The solution was centrifuged at maximum speed and the supernatant was transferred to a fresh tube. Ten μL of the cell lysate were loaded onto a 12.5% polyacrylamide gel for Western blot analysis. The protein was transferred to BA85 Protran nitrocellulose (GE Healthcare) with a semi-dry transfer apparatus (Bio-Rad) at 15 V for 30 min. The nitrocellulose was blocked with 1X PBST (1X PBS, 2% milk, 0.1% Tween-20) for four h. The membrane was incubated overnight at 4 °C with 10 mL PBST and 20 μL α-PurR rabbit serum (polyclonal, 1:500 dilution, Covance). The membrane was washed with 1X PBS before incubating for one hour with 10 mL 1X PBS and 0.5 μL secondary antibody (goat α-rabbit Alexa Fluor 647, Invitrogen). The membrane was washed three times with PBST and three times with PBS prior to imaging on a Licor Odyssey CLx.

### PurR conservation

GenBank files and genome sequences were downloaded from NCBI Assembly for 106 complete reference bacterial genomes. To build the phylogenetic tree, 16S rRNA sequences were extracted from the GenBank files with a custom python script, and three eukaryotic 18S rRNA sequences were included as an outgroup. The rRNA sequences were aligned in MEGA X with ClustalW (Kumar et al., 2018). The best model for tree construction was chosen as General Time Reversible with gamma distributed categories and invariant sites, which had the lowest AIC by MEGA X’s model test software (Nei and Kumar, 2000). The tree was constructed according to this model in MEGA X with extensive grafting and subtree pruning and 750 bootstrap replicates. The tree was modified and annotated in R using the ggtree package (Yu et al., 2017).

Standalone BLAST+ from NCBI was used to determine which reference genomes contained a *B. subtilis* or *B. anthracis* PurR homolog. A BLAST database was created from the FASTA files of the same reference genome sequences used to generate the phylogenetic tree. The 73 N-terminal residues of *B. subtilis* PurR were BLASTed against this database with tblastn and a default E value of 10. This domain is unique to PurR according to Pfam (PF09182). The complete PurR sequence was not used due to homology of the PRT domain with PRT enzymes.

### ChIP-Seq data analysis

Libraries were sequenced on the NextSeq platform at the University of Michigan Advanced Genomics Core. The analysis pipeline read-me is publicly available at GitHub (fill_in). Briefly, quality control was determined with FastQC (https://www.bioinformatics.babraham.ac.uk/projects/fastqc/). Adapters were removed with CutAdapt with arguments --quality-base=33 -a AGATCGGAAGAGC -a CGGAAGAGCACAC -a GGGGGGGGGGGGGGG -n 3 -m 20 --match-read-wildcards -o. Using bowtie2 and arguments -U -q --end-to-end --very-sensitive -p 6 --phred33, wild type *B. subtilis* reads were aligned to the *B. subtilis* NCIB 3610 genome (CP020102) from NCBI Reference Sequence Database. *B. subtilis* (p)ppGpp^0^ reads were aligned to the NCIB 3610 genome with the pBS32 megaplasmid (CP020102 concatenated with CP020103). Using a custom python script, coverage at a 10 bp resolution was bootstrapped and the median coverage of 100 bootstraps was used for downstream analysis. Coverage was smoothed with a custom python script. Enrichment was calculated as the median counts per million in a ChIP sample divided by the median counts per million in its corresponding input sample for each 10 bp window. The input sequencing dataset for the first replicate was low quality, so the input datasets from the same strain background and treatment conditions of the third replicate were used to calculate enrichment for the first replicate.

## ACKNOWLEDGEMENTS

We thank members of the Wang lab for useful discussion and comments on the manuscript. This work is supported by the National Institutes of Health R35 GM127088 from NIGMS (to J.D.W.), R01 AI110740 and R01 AI142400 (to V.T.L.), R21 AI135494 (to M.A.S. and R.G.B.), the Howard Hughes Medical Institute Faculty Scholars Award (to J.D.W.) and National Science Foundation GRFP DGE-1256259 (to B.W.A.).

## Author contributions

B.W.A, V.T.L., M.A.S. R.G.B. and J.D.W. designed the research. B.W.A., M.A.S., J.Y., A.T., H.T., Q.H., V.T.L. and J.D.W. performed experiments. B.W.A., M.A.S., J.Y., V.T.L., R.G.B., and J.D.W. analyzed data. B.W.A. wrote the paper. B.W.A., M.A.S., J.Y., V.T.L., R.G.B., and J.D.W. edited the paper.

## SUPPLEMENTAL FIGURE LEGENDS

**Figure S1.**
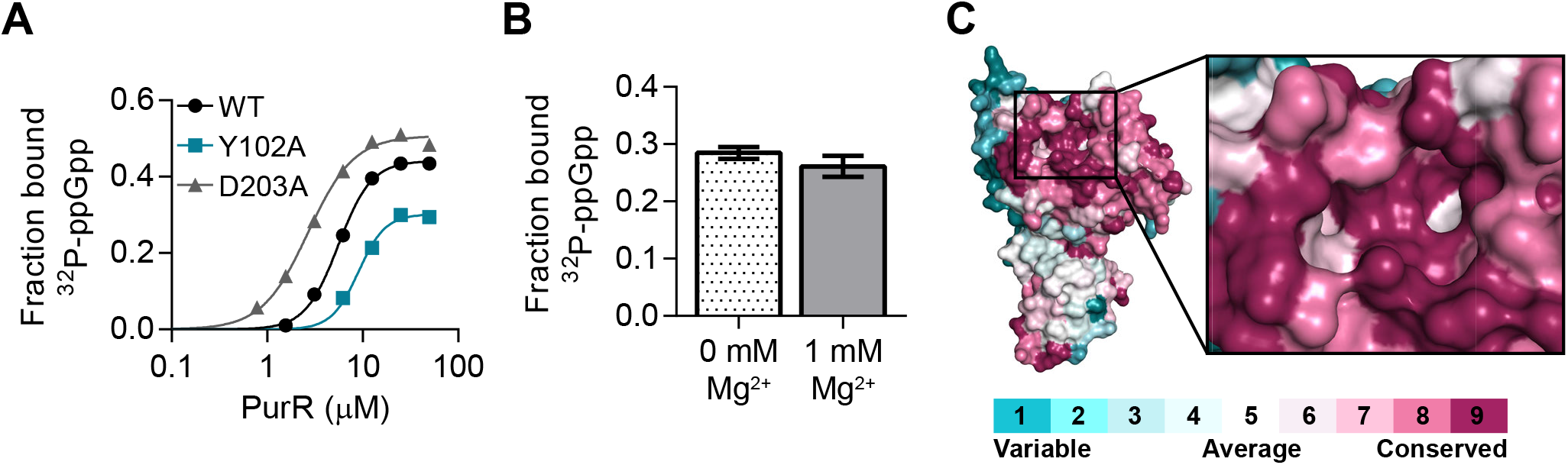
ppGpp binding to *B. subtilis* PurR. **A)** DRaCALA binding curves of ^32^P-ppGpp binding to Y102A and D203A variants of PurR. **B)** Mg^2+^ did not crystallize with PurR-ppGpp, and it is not necessary for the interaction as determined by DRaCALA. **C)** ConSurf analysis of 938 PurR proteins mapped onto PDB ID 1O57. Darker purple indicates more conserved residues. Inset shows the (p)ppGpp binding pocket. Error bars represent SEM of triplicate.

**Figure S2.**
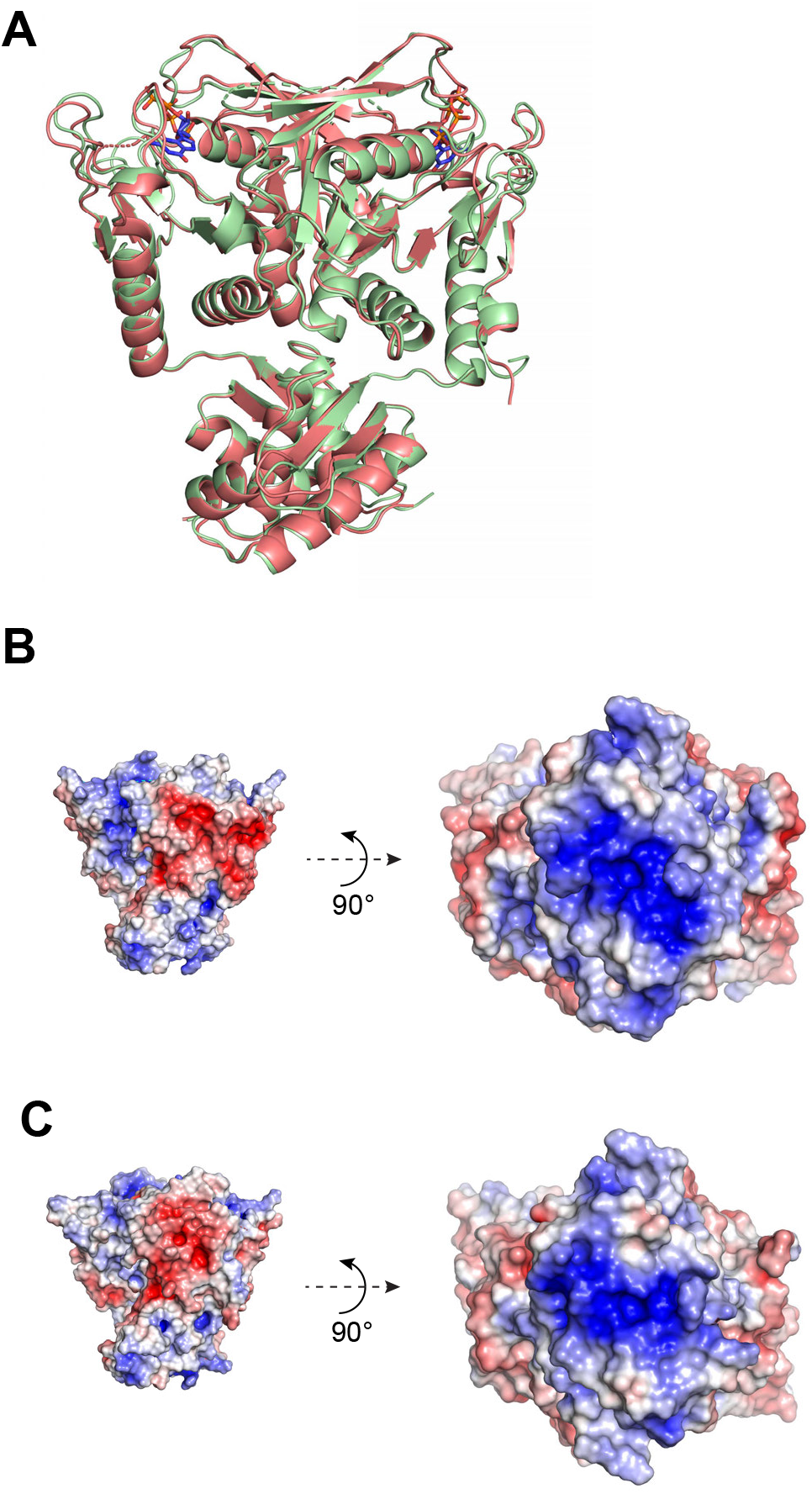
Effect of ppGpp on PurR conformation and electrostatics. **A)** Overlay of *B. subtilis* apo PurR (green; PDB ID 1O57) and ppGpp-bound PurR (salmon) dimer. **B)** Poisson-Boltzmann continuum electrostatics of the DNA binding domain of apo PurR. **C)** Poisson-Boltzmann continuum electrostatics of the DNA binding domain of ppGpp-bound PurR. Scale of electrostatic potential is −5 (red) to +5 (blue) in all figures.

**Figure S3.**
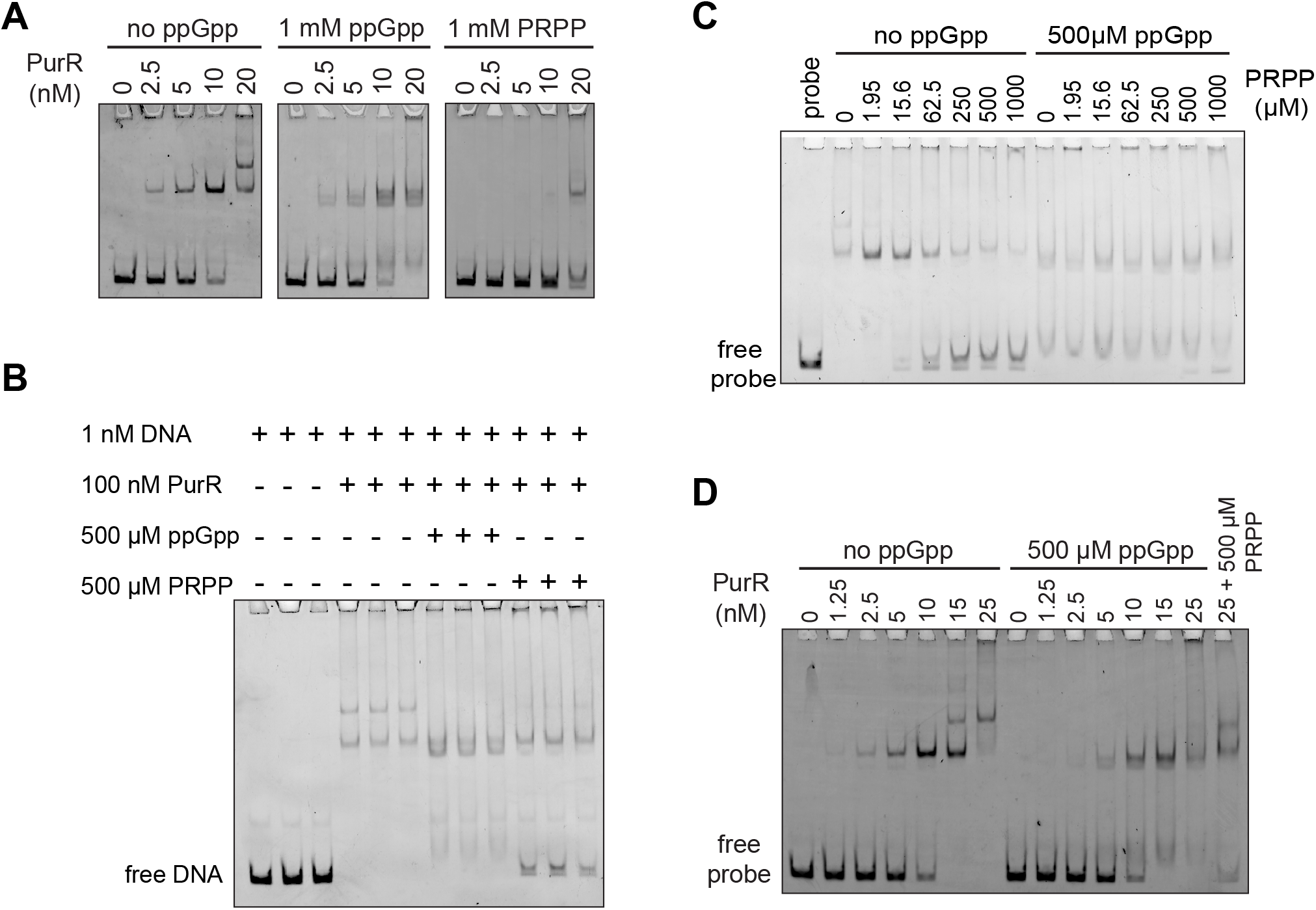
EMSAs of the effect of ppGpp and PRPP on PurR-DNA interaction. **A)** EMSAs showing PurR binding to a 202 bp DNA probe with increasing PurR concentrations in the presence of ppGpp or PRPP. **B)** EMSA showing the individual effects of ppGpp and PRPP on the binding of 100 nM PurR to DNA. **C)** EMSA showing the effect of ppGpp and PRPP competition on PurR binding to a 202 bp DNA probe at a PurR concentration of 100 nM. **D)** EMSA showing the effect of ppGpp on PurR interaction with DNA. DNA fragment used is 202 bp fragment upstream of the *pur* operon transcription start site. These EMSAs were performed at low KCl concentrations and with no nonspecific DNA, and the probe was nonspecifically labeled (see Materials and Methods).

**Figure S4.**
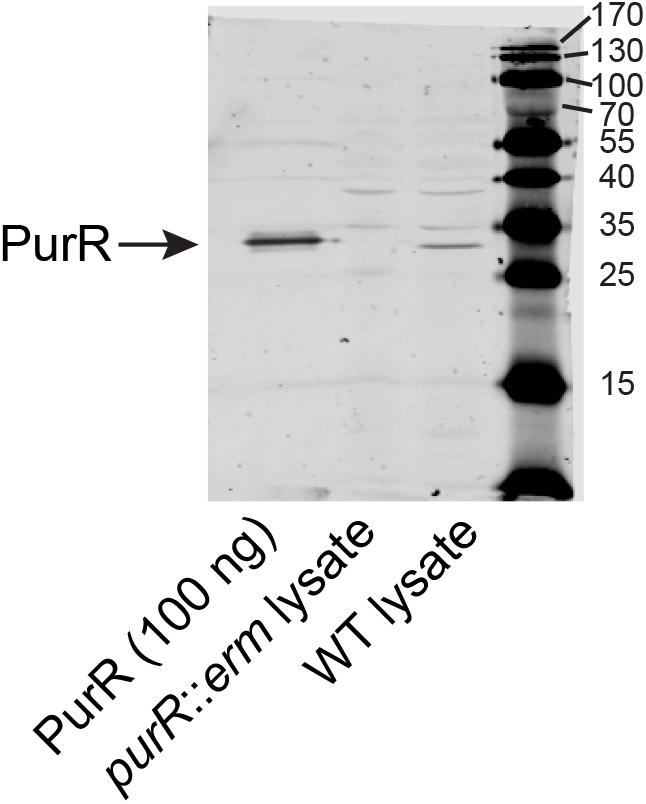
Western blot showing specificity of polyclonal anti-PurR antibody. Western blot of PurR in wild type and *purR::ermHI B. subtilis*. Purified untagged *B. subtilis* PurR included as a control. Molecular weight markers in the ladder refer to kilodaltons.

**Figure S5.**
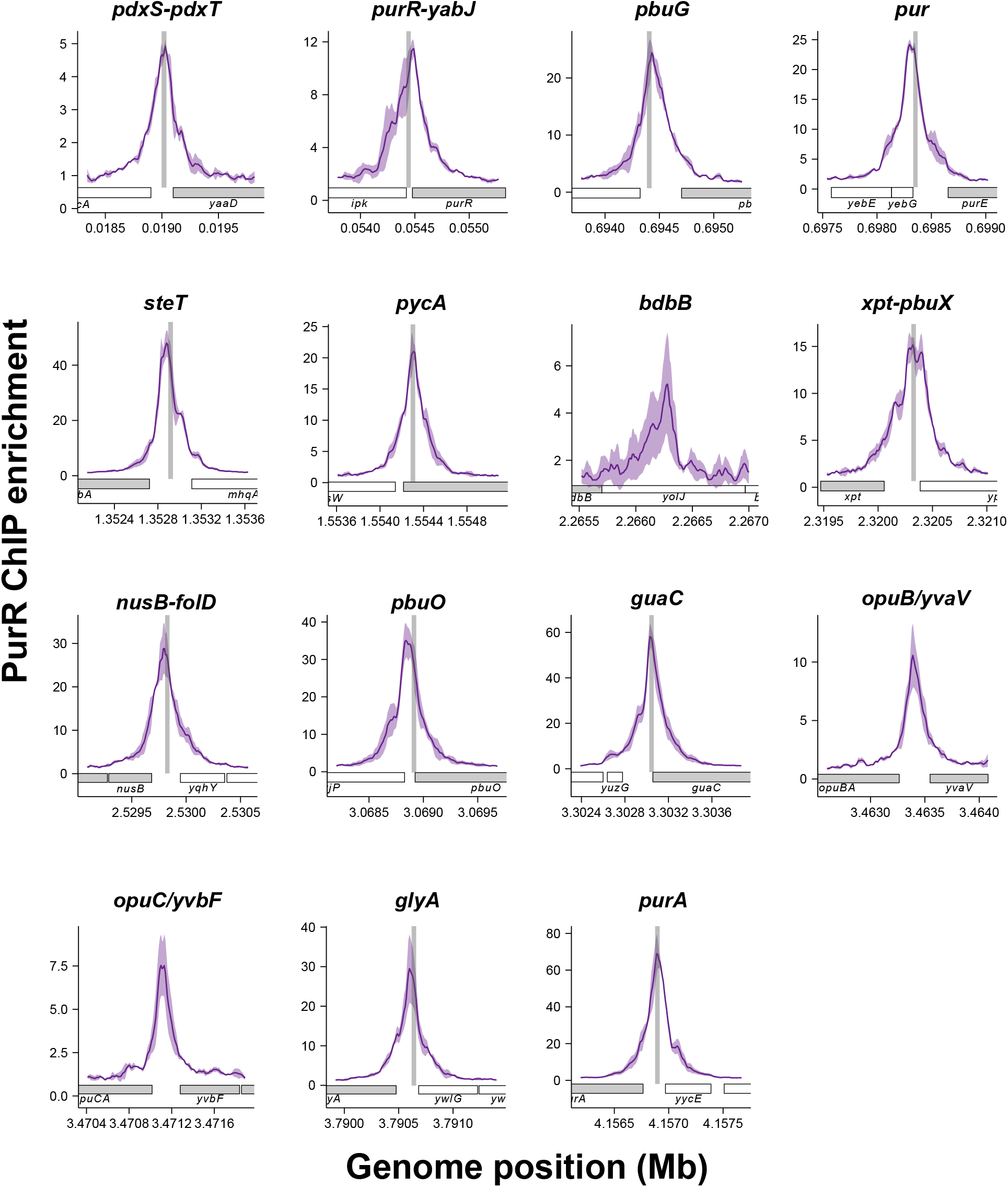
PurR binding sites during growth with nucleobases. The 15 PurR ChIP peaks obtained during growth of *B. subtilis* with ACGU nucleobases. Solid line represents mean of triplicate and shaded region represents standard deviation. Shaded gray vertical box represents the PurBox sequences identified at nine of the PurR binding sites. Genes known or hypothesized to be regulated by PurR are colored gray. See Table 2 for more information.

**Figure S6.**
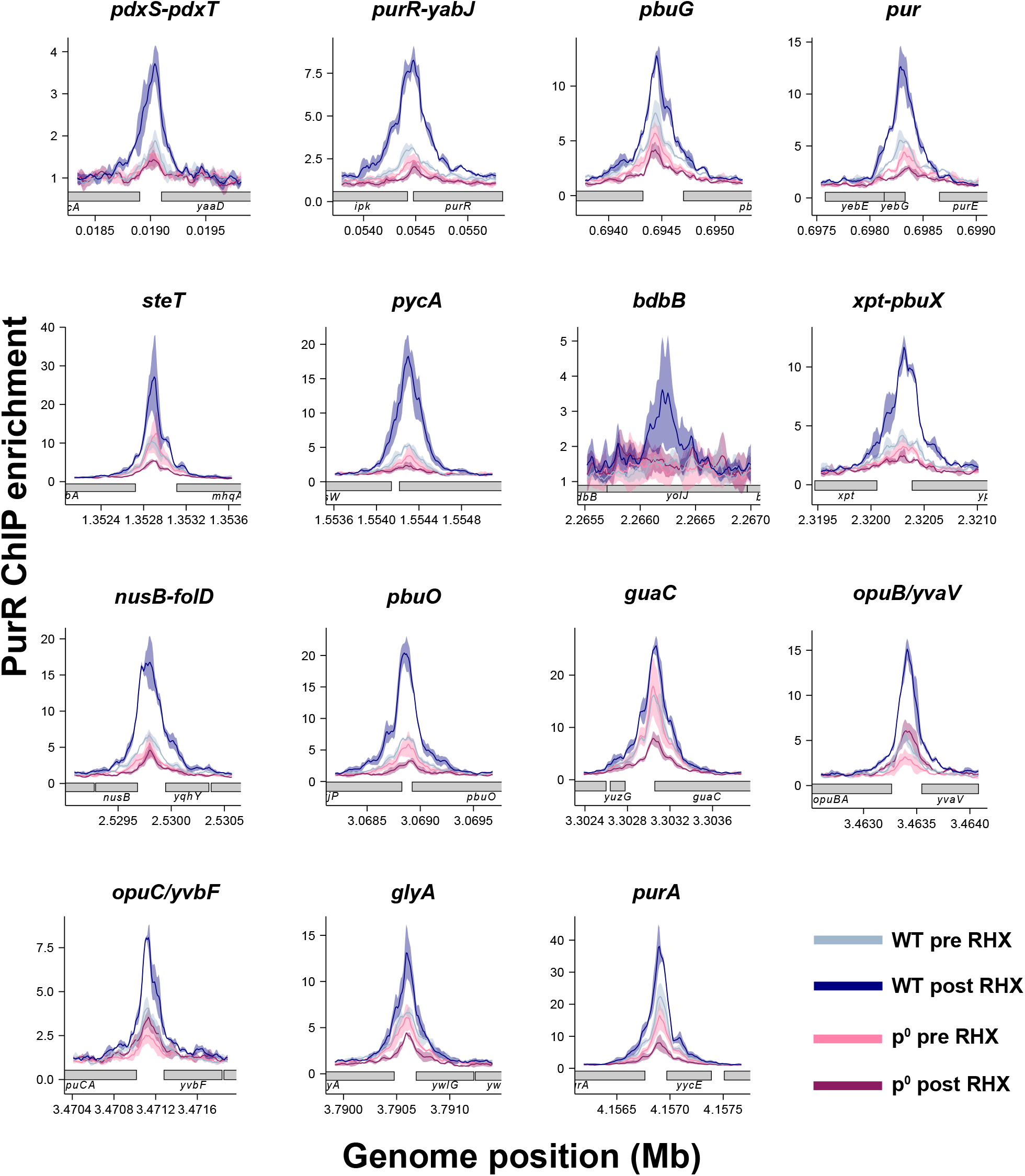
(p)ppGpp induction increases PurR enrichment at all 15 binding sites. The 15 PurR binding sites determined by ChIP-seq before and after (p)ppGpp induction by arginine hydroxamate (RHX). Wild type (WT) before RHX shown in cyan and after RHX shown in blue. (p)ppGpp-null (p^0^) before RHX shown in pink and after RHX shown in maroon. Solid line represents mean of triplicate and shaded region represents standard deviation. See Table 2 for more information.

**Figure S7.**
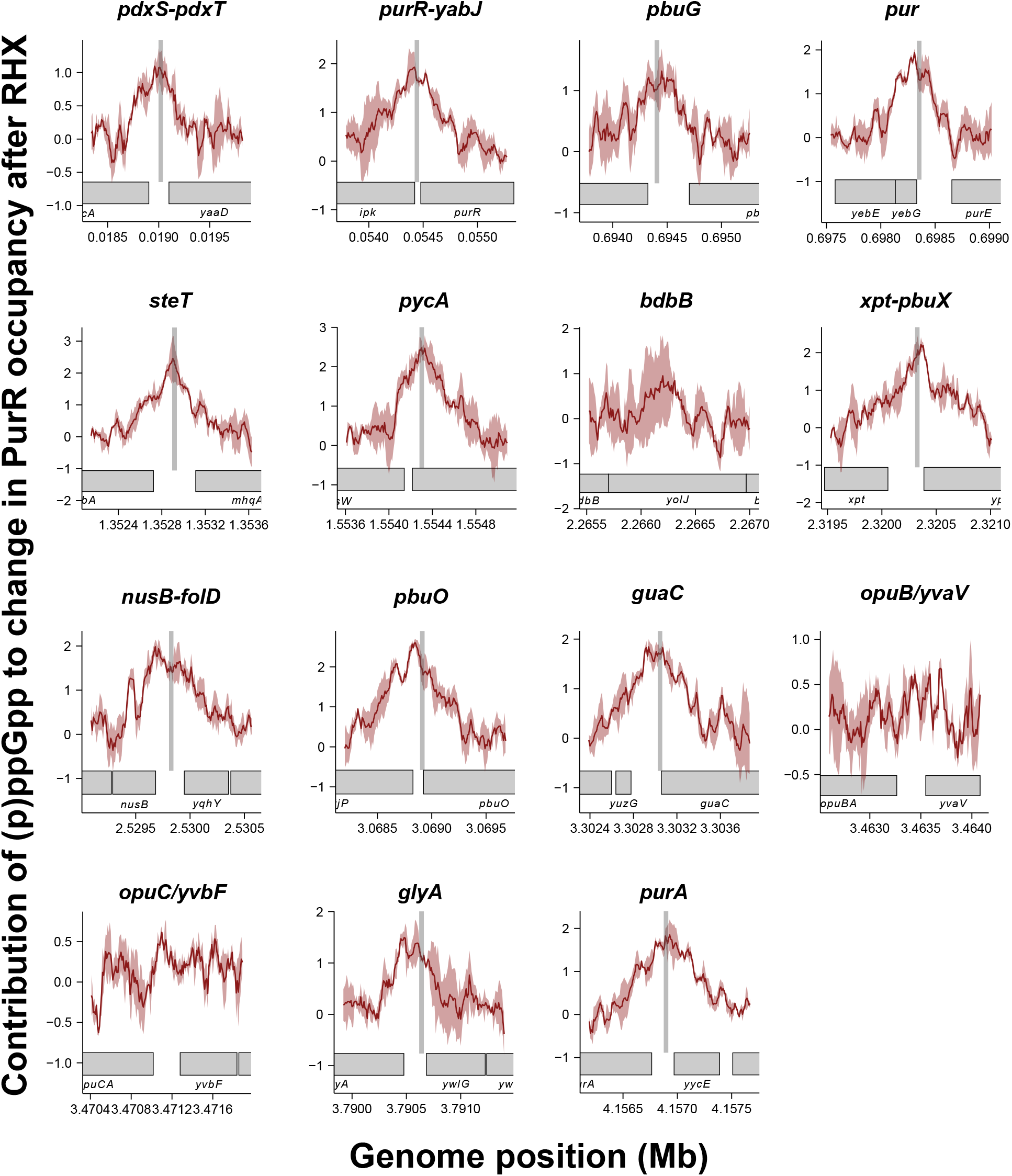
Contribution of (p)ppGpp to change in PurR occupancy. Contribution of (p)ppGpp to change in PurR occupancy for each of the 15 PurR binding sites. The change in PurR occupancy after RHX treatment was calculated for WT and (p)ppGpp^0^, and the ratio of the change in WT over (p)ppGpp^0^ was used as the contribution of (p)ppGpp.

**Figure S8.**
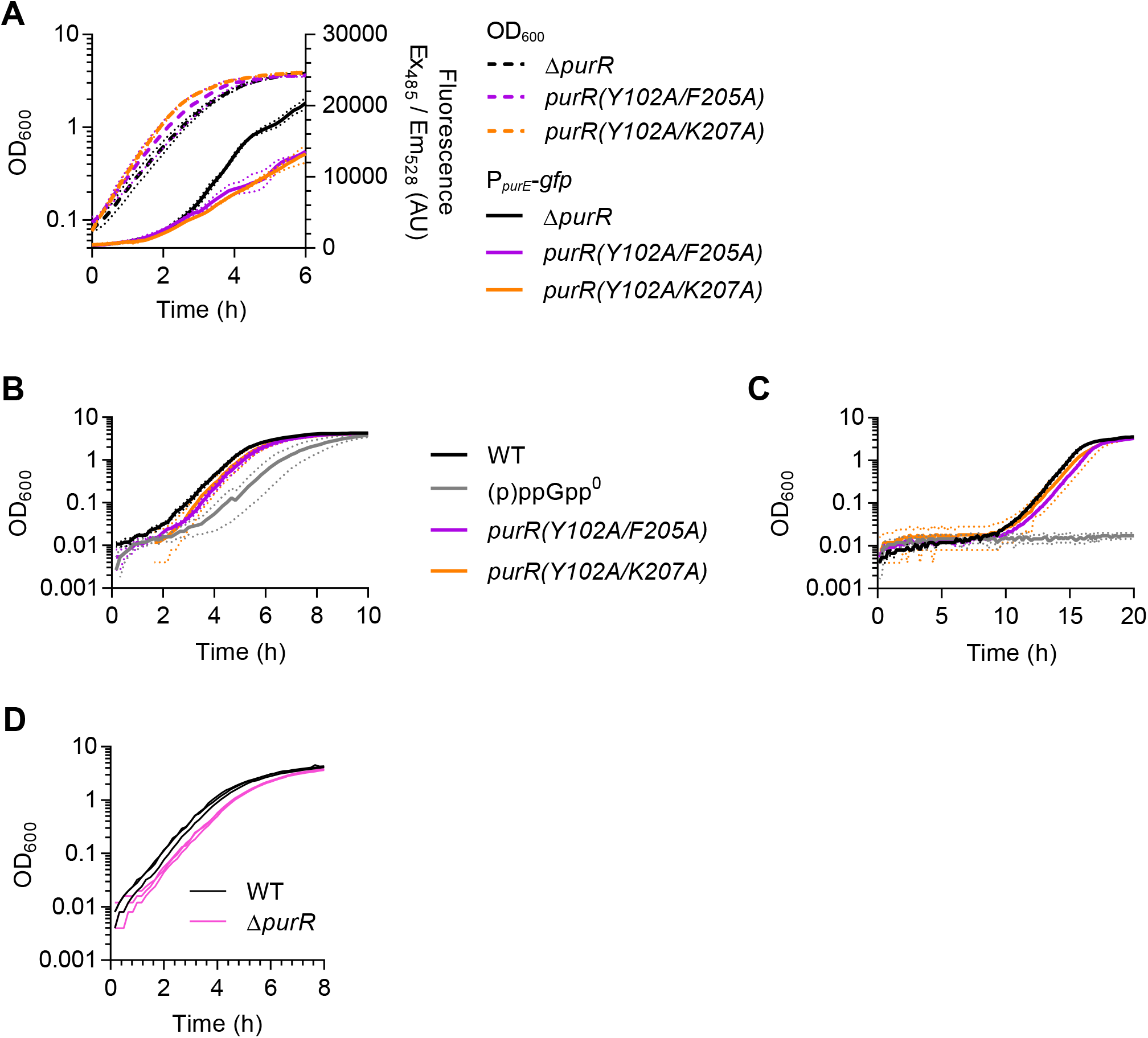
Dysregulated purine synthesis affects nutrient stress adaptation. **A)** Expression of a P*purE*-GFP reporter in Δ*purR*, *purR(Y102A/F205A)*, and *purR(Y102A/K207A) A. subtilis*. Dotted lines represent SEM of triplicate. **B-C)** Same data as shown in Figure 5E-F with log transformation of OD. Cultures were grown in Min+20aa, downshifted to Min, and then outgrown in Min+20aa (C) or Min (D). Error (dotted lines) is SEM of triplicate. **D)** Growth of wild type and ΔpurR in Min+20aa following nutrient downshift from Min+20aa to Min media and subsequent upshift back to Min+20aa. Individual traces of three replicates shown.

## SUPPLEMENTAL TABLES

**Table S1.** pppGpp targets determined by screening ORFeome libraries using DRaCALA.

| Locus ID | Protein | Z-score |
| --- | --- | --- |
| His-tagged library |  |  |
| BA3162 | 5'-nucleotidase, putative GMP -> guanosine | 6.27 |
| BA4322 | 5'-nucleotidase family protein | 5.47 |
| BA4672 | spo0B-associated GTP-binding protein (Obg) | 4.31 |
| BA4455 | membrane protein, putative | 4.23 |
| BA1525 | GTPase family protein (DER. EngA) | 3.97 |
| BA0063 | hypoxanthine phosphoribosyltransferase (hpt1) | 3.92 |
| BA4788 | hypothetical protein 152 aa with heme-binding site, iron transfer operon | 3.52 |
| BA5734 | tRNA modification GTPase TrmE | 3.27 |
| BA4098 | hypothetical protein yuaY | 3.24 |
| BA2853 | cell division protein DivIC | 3.10 |
| Pxo1_47 | hypothetical protein pxo1_47 | 3.05 |
| BA4915 | acetyl-CoA synthetase | 2.98 |
| BA1591 | xanthine phosphoribosyltransferase | 2.83 |
| BA1692 | hypothetical protein | 2.77 |
| BA5074 | hypoxanthine phosphoribosyltransferase (hpt2) | 2.72 |
| BA5716 | adenylosuccinate synthetase purA | 2.69 |
| BA1998 | NH(3)-dependent NAD(+) synthetase (NadE) | 2.55 |
| BA3589 | hypothetical protein | 2.53 |
| HisMBP-tagged library |  |  |
| BA1591 | xanthine phosphoribosyltransferase | 6.18 |
| BA3162 | 5'-nucleotidase, putative GMP -> guanosine | 5.37 |
| BA4672 | spo0B-associated GTP-binding protein (Obg) | 4.95 |
| BA1925 | cobalamin synthesis protein/P47K family protein | 3.87 |
| BA5472 | D-amino acid aminotransferase | 3.79 |
| BA1531 | DNA-binding protein HU | 3.74 |
| BA0692 | hypothetical protein | 3.54 |
| BA1828 | GTP-binding protein HflX | 3.51 |
| BA4699 | transcriptional regulator, MarR family | 3.33 |
| BA0596 | nicotinate phosphoribosyltransferase, putative | 3.32 |
| BA1407 | hypothetical protein | 3.26 |
| BA1212 | hypothetical protein (YjbM) | 3.24 |
| Pxo1_60 | hypothetical protein pxo1_60 | 3.21 |
| BA3140 | hypothetical protein pxo1_60 | 3.15 |
| BA0063 | hypoxanthine phosphoribosyltransferase (hpt1) | 3.05 |
| BA0150 | polysaccharide deacetylase, putative | 2.98 |
| BA2932 | glutathionylspermidine synthase, putative | 2.93 |
| BA0306 | DNA ligase, NAD-dependent | 2.91 |
| BA2254 | hypothetical protein | 2.88 |
| BA4700 | organic hydroperoxide resistance protein | 2.85 |
| BA0880 | hypothetical protein | 2.84 |
| BA2278 | hypothetical protein | 2.83 |
| BA0721 | hypothetical protein | 2.76 |
| BA3131 | alcohol dehydrogenase, zinc- containing | 2.76 |
| BA5251 | hypothetical protein | 2.70 |
| BA2837 | hypothetical protein | 2.68 |
| BA2636 | sensor histidine kinase | 2.60 |
| BA2470 | hypothetical protein | 2.59 |
| BA1349 | hypothetical protein | 2.57 |
| BA4397 | DNA repair protein RecN | 2.57 |
| BA2163 | HD domain protein | 2.55 |
| BA2625 | hypothetical protein | 2.53 |
| BXB0074 | hypothetical protein BXB0074 | 2.52 |
| BA1267 | hypothetical protein | 2.51 |
| BA4807 | hypothetical protein | 2.50 |

**Table S2.**
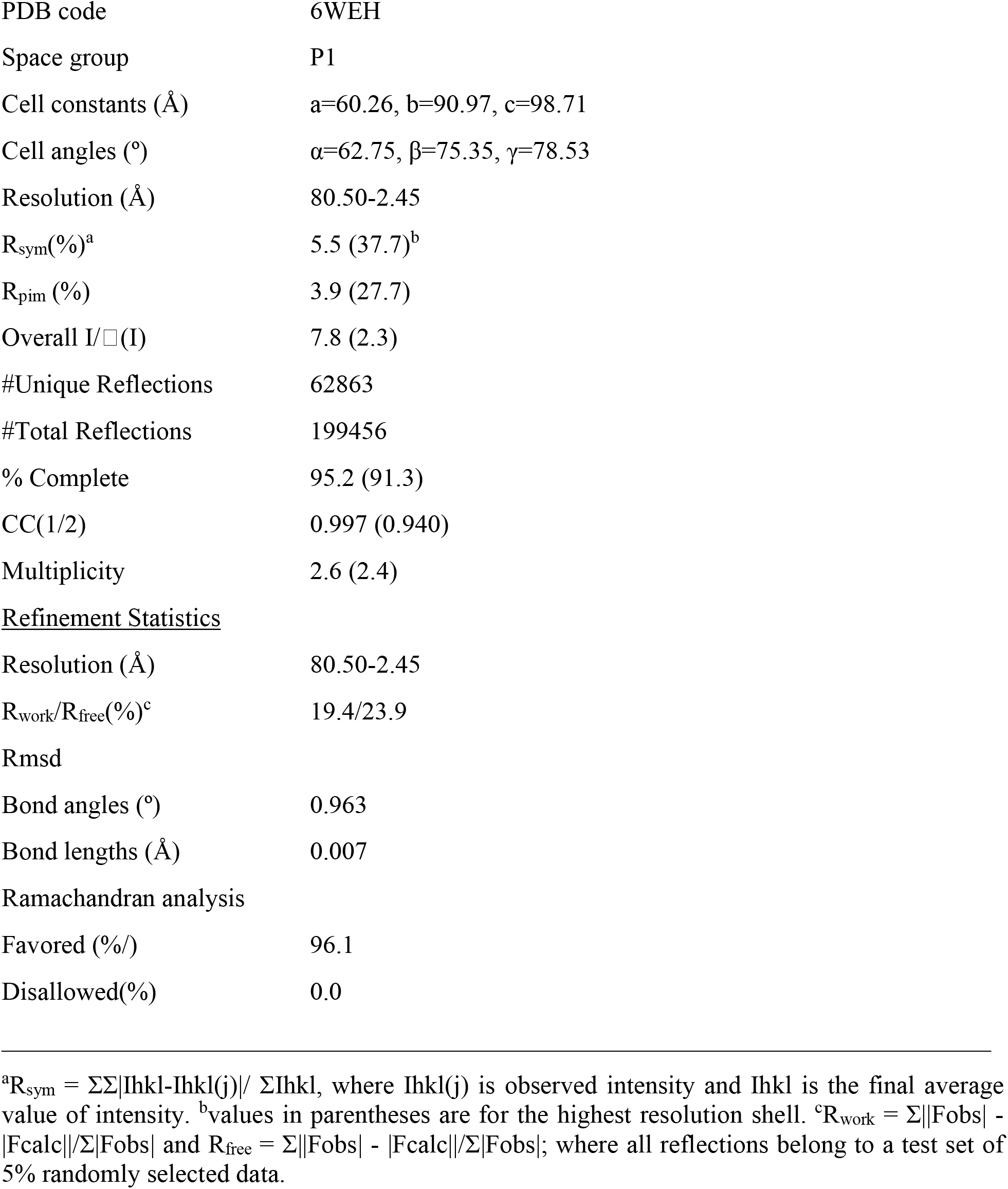
Data collection and refinement statistics for *B. subtilis* PurR-ppGpp structure.

**Table S3.**
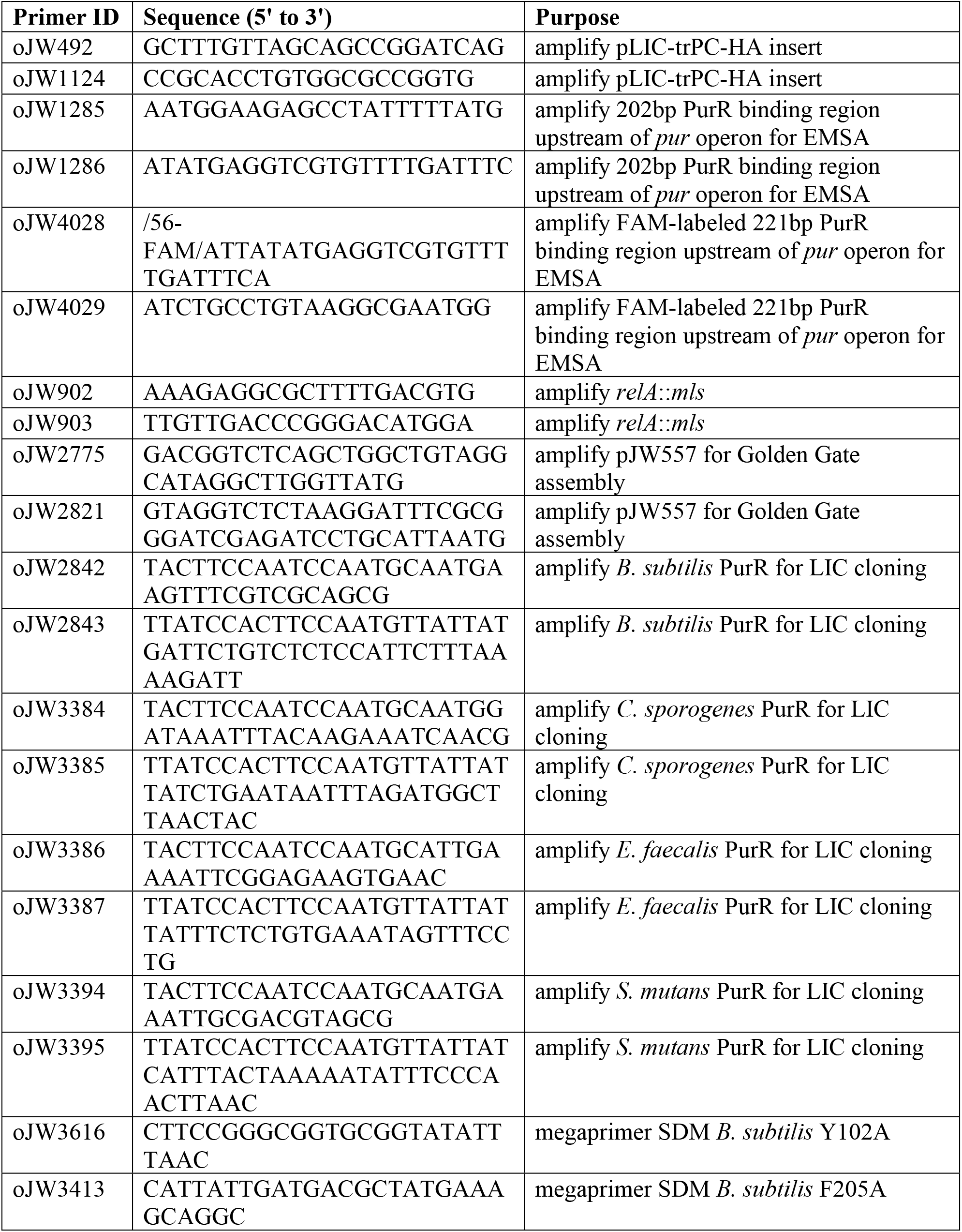
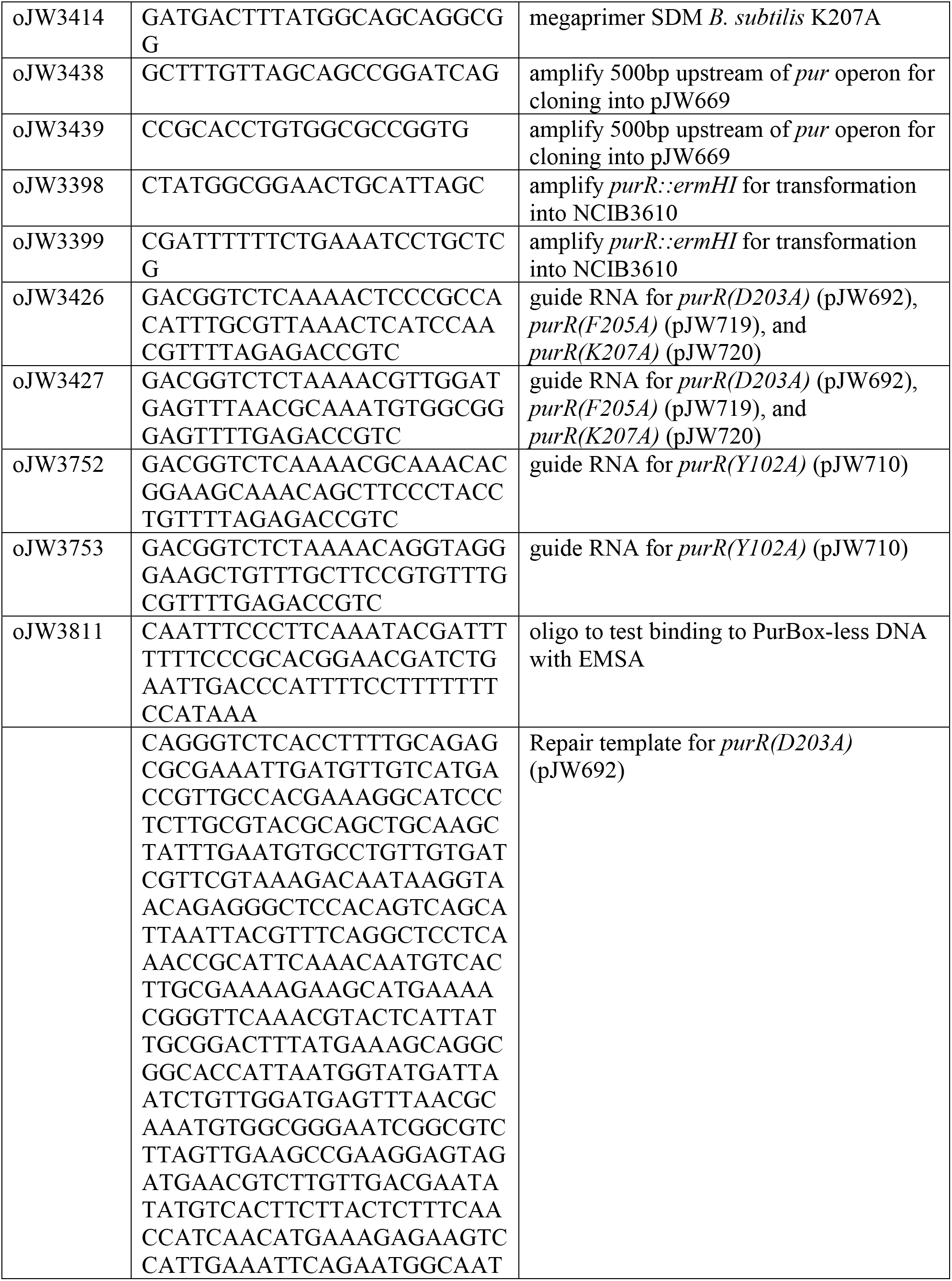
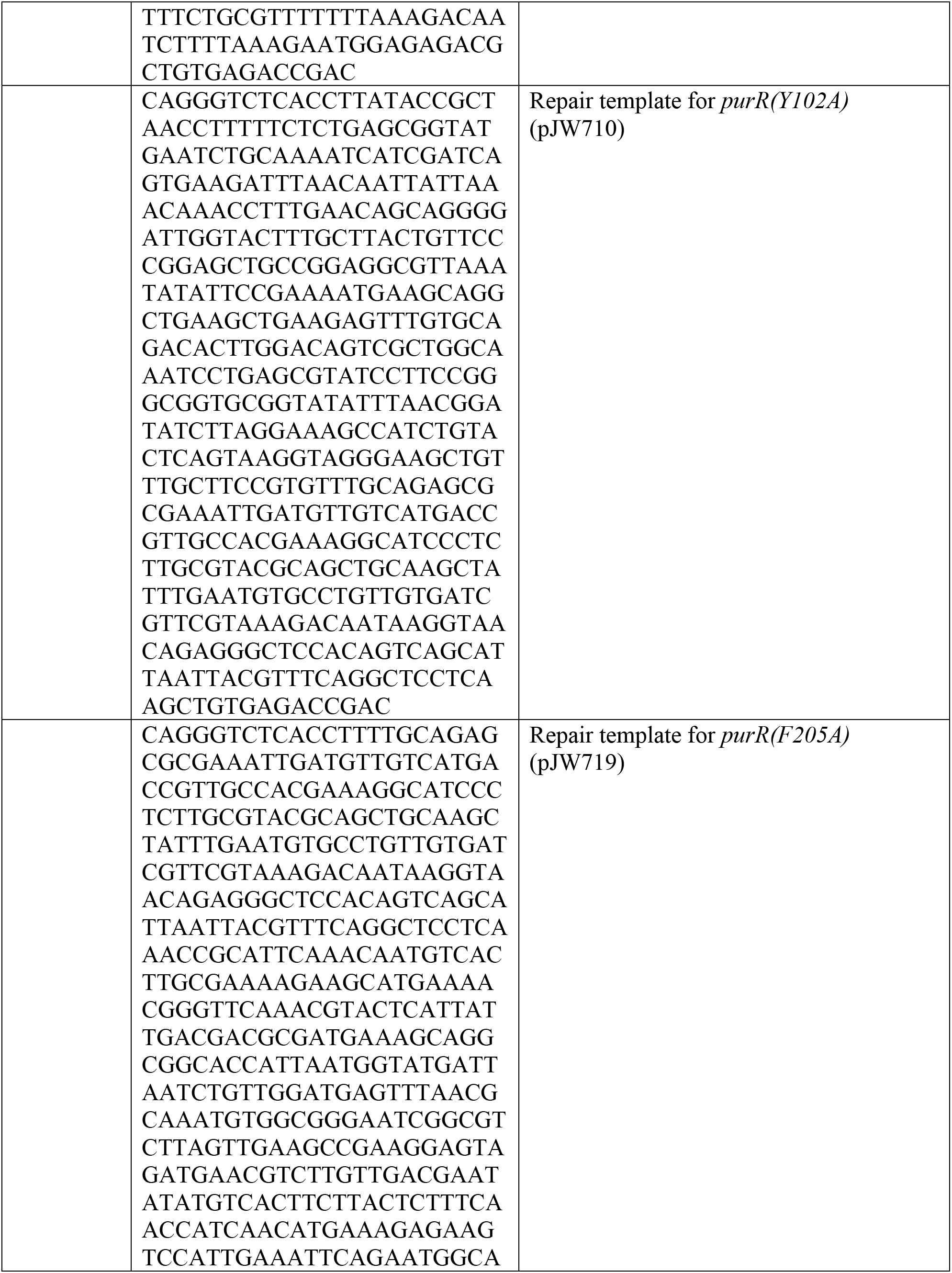
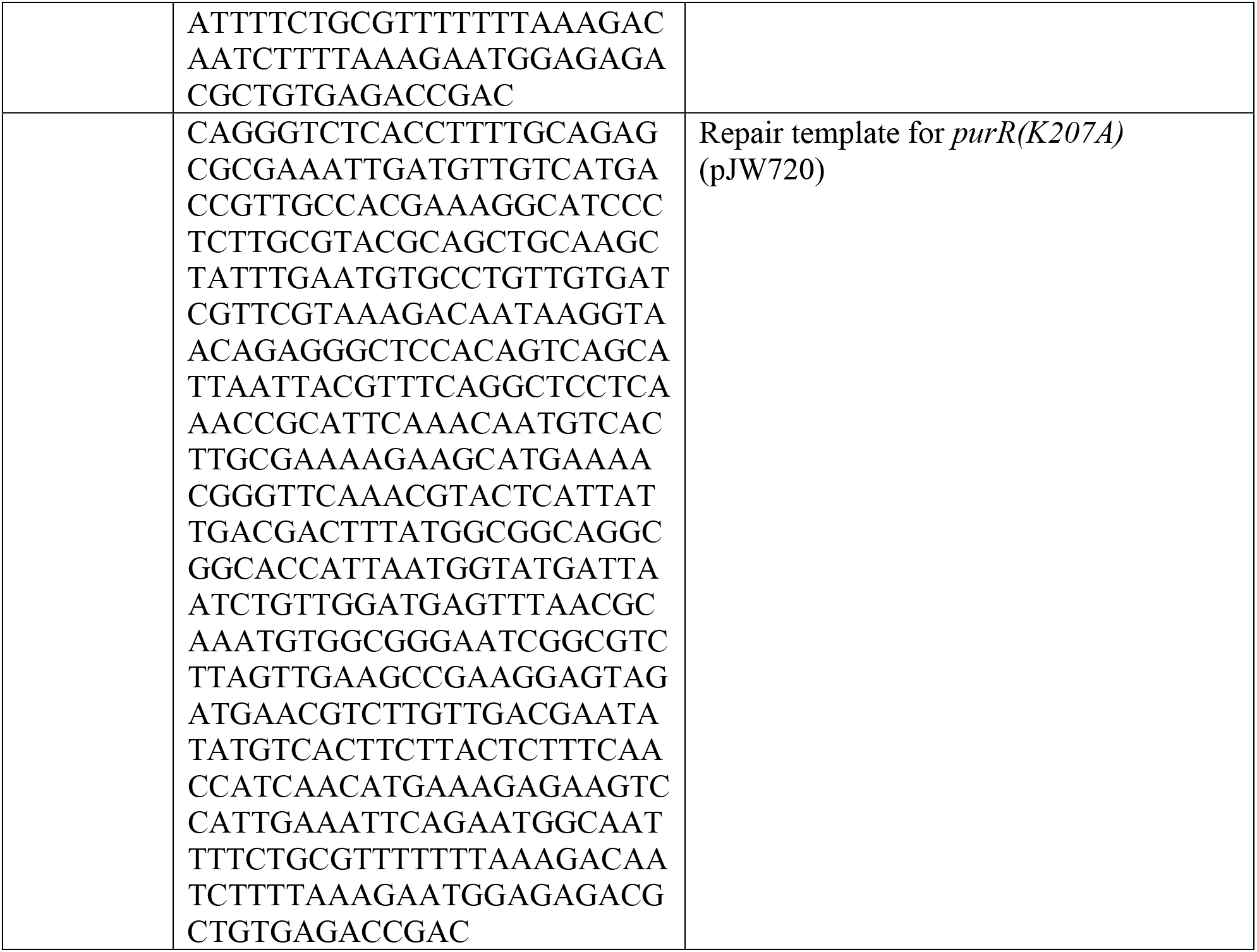
List of primers.

**Table S4.** List of plasmids.

| Plasmid ID | Construct | Purpose | Reference |
| --- | --- | --- | --- |
| pJW270 | pLIC-trPC-HMA | Expression vector | Stols et al., 2002 |
| pJW664 | pLIC-trPC-HMA *B. subtilis* *purR* | Expression of HisMBP-tagged *B. subtilis* PurR | this work |
| pJW669 | pPB41 with promoterless superfolder GFP | Cloning promoter of interest controlling sfGFP expression | this work |
| pJW686 | pJW669 P<sub>*purE*</sub>-sfGFP | GFP reporter for P<sub>*purE*</sub> operon, for recombination at *amyE* | this work |
| pJW692 | pPB41 with *purR*(D203A) gRNA and repair template | CRISPR recombineering *purR*(D203A) | this work |
| pJW710 | pPB41 with *purR*(Y102A) gRNA and repair template | CRISPR recombineering *purR*(Y102A) | this work |
| pJW719 | pPB41 with *purR(F205A)* gRNA and repair template | CRISPR recombineering *purR(F205A)* | this work |
| pJW720 | pPB41 with *purR(K207A)* gRNA and repair template | CRISPR recombineering *purR(K207A)* | this work |
| pJW558 | pDR110 with P<sub>*ilvB*</sub>-GFPmut2<sup>A206K</sup> | P<sub>*ilvB*</sub>-GFPmut2<sup>A206K</sup> reporter | Fung et al., 2020 |
| pVL791 | pET19-based expression vector with N-terminal His<sub>6</sub> tag | Expression of His-tagged ORFeome library | Roelofs et al., 2011 |
| pVL847 | pET19-based expression vector with N-terminal His<sub>6</sub>-MBP tag | Expression of HisMBP-tagged ORFeome library | Lee et al., 2007 |

**Table S5.**
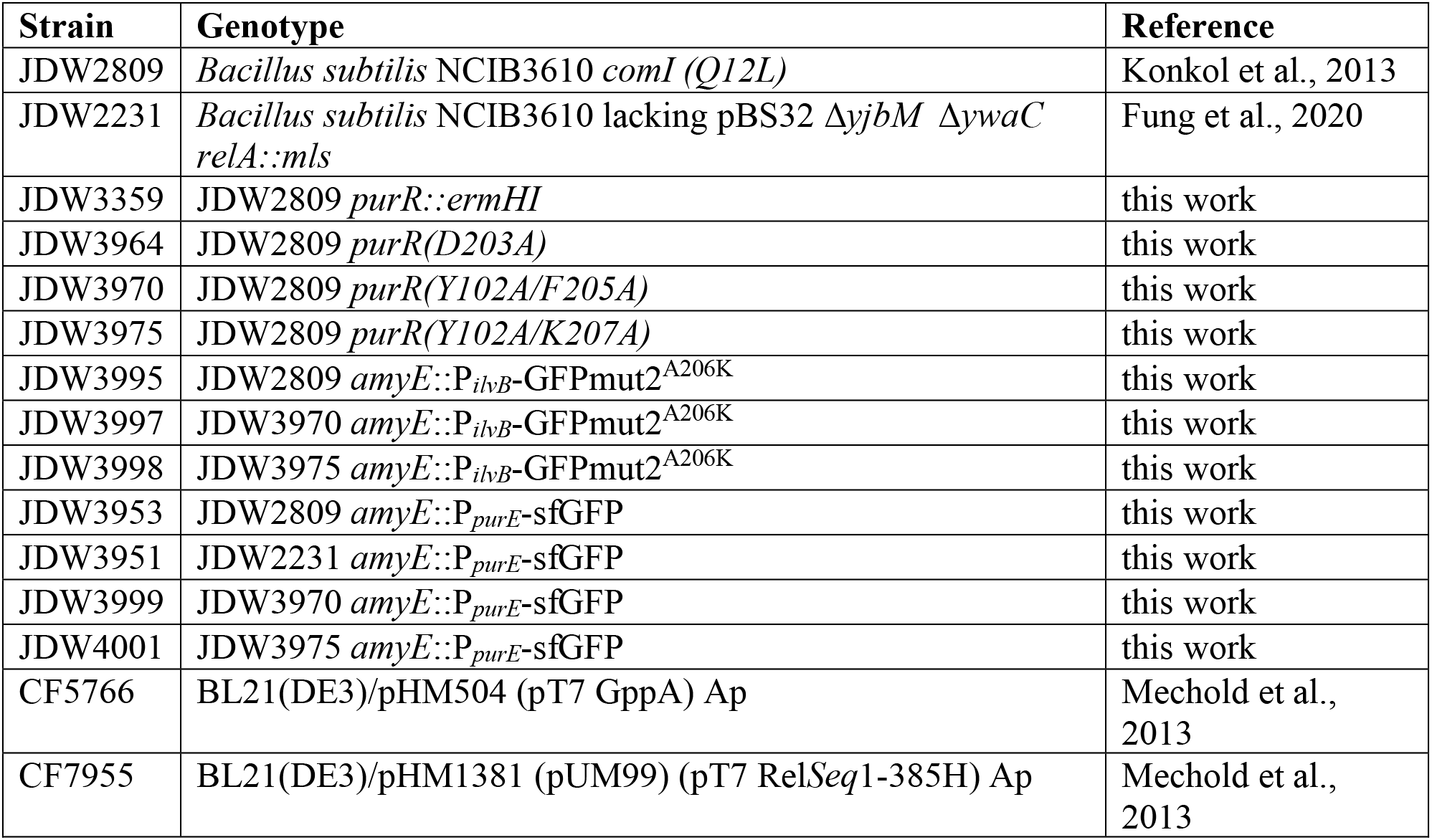
List of strains.

## Notes

### Competing Interest Statement

The authors have declared no competing interest.

